# Differential requirement for centriolar satellites in cilium formation among different vertebrate cells

**DOI:** 10.1101/478974

**Authors:** Ezgi Odabasi, Signe K. Ohlsen, Seref Gul, Ibrahim H. Kavakli, Jens S. Andersen, Elif N. Firat-Karalar

## Abstract

Centriolar satellites are ubiquitous in vertebrate cells. They have recently emerged as key regulators of centrosome/cilium biogenesis, and their mutations are linked to ciliopathies. However, their precise functions and mechanisms of action, which potentially differ between cell types, remain poorly understood. Here, we generated retinal pigmental and kidney epithelial cells lacking satellites by genetically ablating PCM1 to investigate their functions. While satellites were essential for cilium assembly in retinal epithelial cells, kidney epithelial cells lacking satellites still formed full-length cilia but at significantly lower levels, with reduced centrosomal levels of key ciliogenesis factors. Using these cells, we identified the first satellite-specific functions at cilia, specifically in regulating ciliary content, Hedgehog signalling, and epithelial cell organization. However, other satellite-linked functions, namely proliferation, cell cycle progression and centriole duplication, were unaffected in these cells. Quantitative transcriptomic and proteomic profiling revealed that loss of satellites scarcely affects transcription, but significantly alters the proteome, particularly actin cytoskeleton pathways and neuronal functions. Together, our findings identify cell type-specific roles for satellites and provide insight into the phenotypic heterogeneity of ciliopathies.

## Introduction

The vertebrate centrosome/cilium-complex is composed of centrosomes, cilia and centriolar satellites. The evolutionarily conserved microtubule-based cylindirical structures, centrioles, recruit the pericentriolar material to form the centrosome (Carvalho-Santos et al, 2011; Hodges et al, 2010). In quiescent and differentiated cells, the older mother centriole functions as a basal body to nucleate the formation of flagella and cilia, microtubule-based structures projecting out from the apical surface of cells. In addition to these conserved structures, specific to vertebrate cells is the array of 70-100 nm membrane-less granules termed centriolar satellites, which localize to and move around the centrosome/cilium-complex in a microtubule-and dynein/dynactin-dependent manner and are implicated in active transport or sequestration of centrosome and cilium proteins (Hori & Toda, 2017; Kubo et al, 1999; Tollenaere et al, 2015).

The assembly and function of the centrosome/cilium-complex is regulated in response to cell-cycle cues and environmental factors. In interphase cells, the centrosome nucleates and organizes the microtubule array, which functions in vesicular trafficking, cell motility and maintaining cell shape and polarity (Luders & Stearns, 2007; Paz & Luders, 2018). During mitosis, centrosomes duplicate and form the bipolar mitotic spindle (Nigg et al, 2014; Nigg & Holland, 2018). In quiescent cells, centrioles dock to the plasma membrane to form the primary cilia, which function as a nexus for developmentally important signaling pathways like Hedgehog and Wnt signaling (Nachury, 2014). Primary cilium formation is a highly complex and regulated process that is mediated through intracellular pathway in the majority of cells and extracellular pathway in polarized epithelial cells (Sorokin, 1962; Sorokin, 1968). Given that centrosomes and cilia are indispensable for key cellular processes, their structural and functional defects are associated with a variety of human diseases including cancer and ciliopathies (Bettencourt-Dias et al, 2011; Nigg et al, 2014). Elucidating the mechanisms that regulate the assembly and function of the centrosome/cilium complex in time and space is required to define the molecular defects underlying these diseases.

Centriolar satellites have recently emerged as key regulators of centrosome/cilium-complex biogenesis (Barenz et al, 2011; Hori & Toda, 2017; Tollenaere et al, 2015). Consistently, mutations affecting satellites components are linked to diseases associated with defects in the centrosome/cilium complex such as ciliopathies, primary microcephaly and schizophrenia (Bradshaw & Porteous, 2012; Hori & Toda, 2017; Kodani et al, 2015; Tabares-Seisdedos & Rubenstein, 2009). Ciliopathies are genetic diseases characterized by a multitude of symptoms including retinal degeneration and polycystic kidney disease (Braun & Hildebrandt, 2017; Reiter & Leroux, 2017). Why defects of the broadly expressed centrosome/cilium complex components are reflected as clinically restricted and heterogeneous phenotypes among different tissue types is unknown. Corroborating the link between satellites and ciliopathies, loss of satellites in zebrafish caused characteristics of cilium dysfunction, analogous to those observed in human ciliopathies (Stowe et al, 2012).

Although centriolar satellites are ubiquitous in vertebrate cells, their size and number vary among different cell types, and change in response to signals including cell cycle cues and stress (Nielsen et al, 2018; Rai et al, 2018; Villumsen et al, 2013). In different cell types and tissues, their cellular distribution ranges from clustering at the centrosomes, nuclear envelope and/or basal bodies, to scattering throughout the cytoplasm (Kubo & Tsukita, 2003; Srsen et al, 2009; Vladar & Stearns, 2007). This variation suggests that satellites might have cell type and tissue-specific functions. Elucidation of these functions could provide important insight into the mechanisms underlying the phenotypic heterogeneity of ciliopathies.

Over 100 proteins have been defined as satellite components through their co-localization with or proximity to PCM1 (pericentriolar material-1), the scaffolding protein of satellites that mediates their assembly and maintenance (Gupta et al, 2015; Hori & Toda, 2017). Phenotypic characterization of various satellite proteins has defined functions in cilium formation, centrosome duplication, microtubule organization, mitotic spindle formation, chromosome segregation, actin filament nucleation and organization, stress response and autophagy (Firat-Karalar et al, 2014; Hori & Toda, 2017; Joachim et al, 2017; Kim et al, 2012; Kodani et al, 2015; Pampliega et al, 2013; Staples et al, 2014; Tang et al, 2013; Villumsen et al, 2013; Wang et al, 2016). However, these may not be satellite-specific functions per se, because all satellite proteins identified so far except for PCM1 localize both to satellites and centrosomes and/or cilia.

Satellite-specific functions have been identified through transient or constitutive depletion of PCM1 from cells, which causes satellite disassembly (Dammermann & Merdes, 2002; Wang et al, 2016). Transient PCM1 depletion causes defects in protein targeting to the centrosome, interphase microtubule network organization, cell cycle progression, cell proliferation, and cilium formation, as well as neuronal progenitor cell proliferation and migration during cortical development (Dammermann & Merdes, 2002; Ge et al, 2010; Kim et al, 2008; Nachury et al, 2007; Zhang et al, 2016). A recent study that generated human PCM1-/-retinal pigmental epithelial (RPE1) cells showed that satellites are essential for cilium assembly, where they sequester the E3 ubiquitin ligase Mib1 and prevent degradation of the key ciliogenesis factor Talpid3, which is required for the recruitment of ciliary vesicles (Wang et al, 2016). However, because RPE1::PCM1-/-cells do not ciliate, they did not allow assessment of satellite-specific functions in ciliary signalling and ciliary targeting of proteins, which are among the predominant phenotypic defects underlying cilipathies. Therefore, a major unresolved question that pertains to our understanding of satellites and their relationship to specific ciliopathies is the identification of the full repertoire of satellite-specific functions, which might vary in different cell types and tissues.

In this study, we generated vertebrate cells that are null for centriolar satellites by ablating the core satellite component PCM1 using genome editing in two different cell types, and studied the cellular and molecular consequences specifically of satellite loss on both centrosome- and cilium-related cellular processes. We chose mouse inner medullary collecting duct cells (IMCD3) cells to complement RPE1 cells because, in contrast to the RPE1 cells, they are Hedgehog responsive, form epithelial spheroids when grown in a three-dimensional matrix, which is dependent on cilia functions, and they ciliate using the extracellular ciliogenesis pathway, whereas RPE1 cells use the intracellular pathway. Together, our results identify satellites as key regulators of ciliogenesis, ciliary signalling, tissue architecture and cellular proteostasis, and provide insight into the mechanisms underlying specific ciliopathi.

## Results

### Disruption of *PCM1* causes loss of centriolar satellites in vertebrate cells

To determine the cellular functions of centriolar satellites and to investigate whether loss of satellites has varying effects on these processes across different cell types, we generated satellite-less cells by disrupting the *PCM1* gene in retinal epithelial RPE1 and kidney epithelial IMCD3 cells. Homozygous null mutations in both alleles of the *PCM1* locus were made using CRISPR/Cas9-mediated genome editing with guides designed to target exon 3 (protein-coding exon 2) in RPE1 and IMCD3 cells. We isolated three PCM1-/-IMCD3 and two PCM1-/-RPE1 clones (hereafter IMCD3 PCM1 KO and RPE1 PCM1 KO) and one control colony (hereafter WT) that was transfected with the plasmid encoding the scrambled gRNA. Sequencing of the PCM1 alleles identified these clones as compound heterozygotes bearing premature stop codons that result from small deletions of less than 20 base pairs and/or insertion of one or two base pairs around the cut site (Figure 1A, S1A-C). Immunoblot analysis of whole-cell lysates showed that PCM1 was not expressed in the IMCD3 and RPE1 PCM1 KO clones (Figure 1B, S1E). Immunofluorescence analysis of these clones relative to controls with two different polyclonal antibodies, one directed against the N-terminal 1-254 amino acids and the other against the C-terminal 1665-2026 amino acids of PCM1, further validated lack of PCM1 expression (Figure 1C, S1D, F). The absence of PCM1 signal in the PCM1 KO clones with the C-terminal PCM1 antibody eliminated the possibility that in-frame gene products downstream of the gRNA target site were initiated, and showed that PCM1 alleles in these clones are likely to be null mutations, which was confirmed by mass-spectrometry based quantitative global proteome analysis described below.

**Figure 1.**
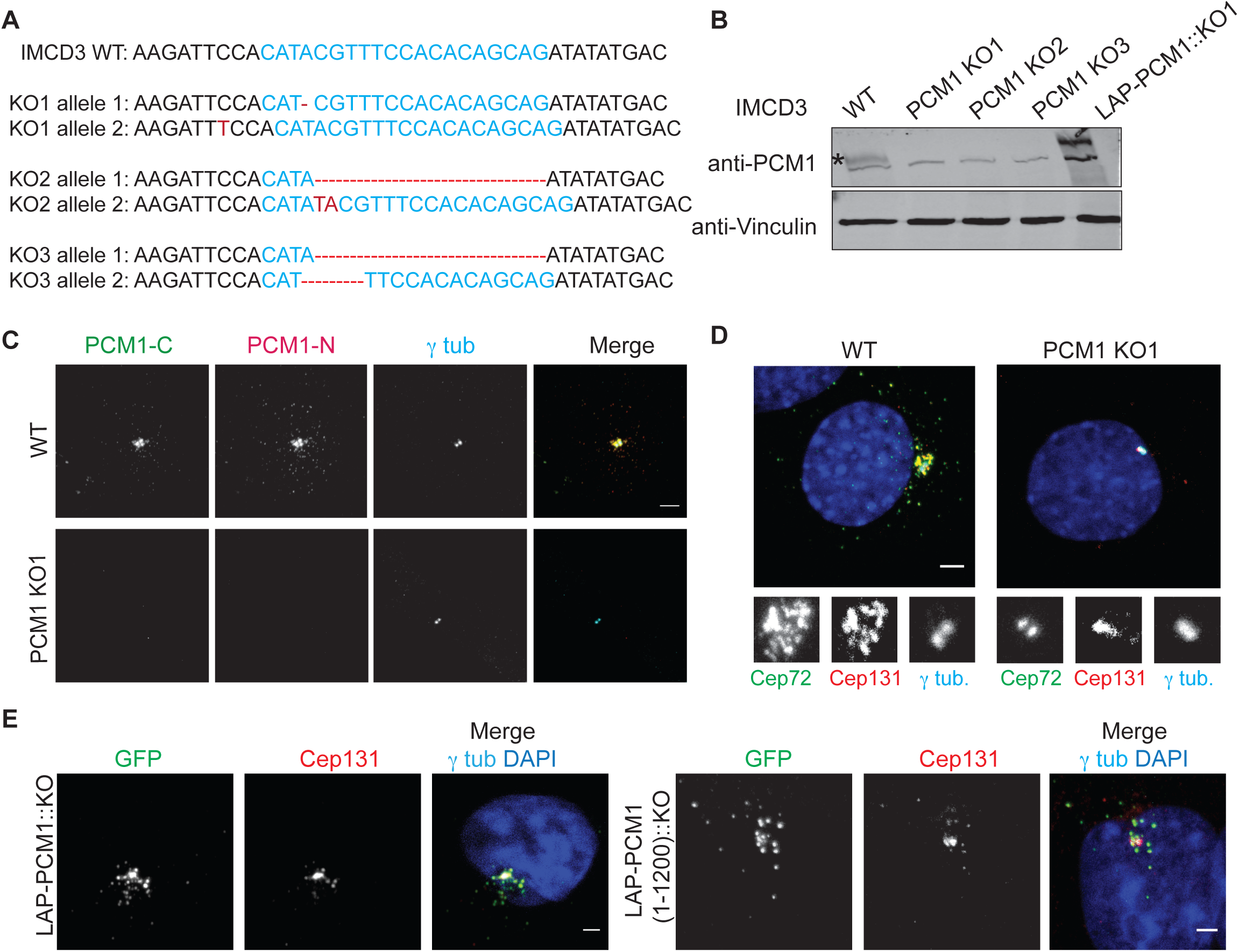
IMCD3 PCM1 KO cells are devoid of satellite structures. **(A)** IMCD3 PCM1 KO clones are all compound heterozygotes with mutations that lead to early stop codons. 1000 bp region around the gRNA-target site was PCR amplified and cloned. Sequencing of five different clones for each line identified one nucleotide (nt) deletion on one allele and one nt insertion on the other for line 1, 16 nt deletion on one allele and 2 nt insertion on the other for line 2 and 16 nt deletion for one allele and 4 nt deletion on the other for line 3. **(B, C)** IMCD3 PCM1 KO cells do not express PCM1. **(B)** Immunoblot analysis of whole-cell lysates from control cells, three IMCD3 PCM1 KO lines and one IMCD3 PCM1 KO::LAP-PCM1 rescue line with PCM1 antibody raised against its N-terminus. **(C)** Immunofluorescence analysis of control and IMCD3 PCM1 KO1 cells. Cells were fixed and stained for centrosomes with anti-³-tubulin antibody and PCM 1 with PCM1-N antibody raised against its N terminal (1-254 amino acids) fragment and PCM1-C antibody raised against its C terminal (1665-2026 amino acids) fragment. Scale bar, 10 μm. **(D)** Localization of satellite proteins Cep72 and Cep131 is restricted to the centrosome in IMCD3 PCM1 KO cells. Cells were fixed and stained for Cep72, Cep131 and gamma-tubulin. DNA was stained with DAPI. Scale bar, 10 μm. **(E)** Stable expression of LAP-PCM1 and LAP-PCM1 (1-3600) rescues the satellite assembly defects in IMCD3 PCM1 KO cells. Cells were fixed and stained with GFP, Cep131 and gamma-tubulin antibodies. DNA was stained with DAPI. Asterisk (*) indicates the band corresponding to full-length PCM1. Scale bar, 10 μm

To confirm that PCM1 KO cells lack satellite structures, we determined the localization of other known satellite components in these cells. While Cep131 and Cep72 localized both to the centrosome and satellites in control cells, their localization was restricted to the centrosomes in IMCD3 PCM1 KO cells, and there were no granules around the centrosome that were positive for satellite markers in these cells (Fig. 1D). Stable expression of LAP-tagged human full-length PCM1 and PCM1 (1-1200) in IMCD3 PCM1 KO clone#1 rescued satellite assembly and localization defects, confirming the specificity of these phenotypes (Fig. 1E). Of note, even though PCM1 (1-1200) expression formed PCM1-positive granules that recruited other satellite components including Cep131, the size of the granules were larger than the ones formed through expression of full-length PCM1 (Fig. 1E), suggesting a regulatory role for the C-terminal part of PCM1 in satellite assembly. These results show that IMCD3 PCM1 KO clones are devoid of satellites and that PCM1 is essential for the assembly of satellites and sequestration of centrosome proteins to the satellites.

### Satellites are required for efficient ciliogenesis, but not for cell proliferation, cell cycle progression and centriole duplication

Many satellite proteins including PCM1 have previously been implicated in ciliogenesis-related functions (Hoang-Minh et al, 2016; Kim et al, 2008; Nachury et al, 2007; Wang et al, 2016). However, the function of satellites during primary cilium formation in different cell types has not been studied in clean genetic backgrounds. To address this, IMCD3 and RPE1 PCM1 KO cells were serum starved for 24 h and 48 h and the percentage of ciliated cells were quantified by staining cells for Arl13b, a marker for the ciliary membrane, and acetylated tubulin, a marker for the ciliary axoneme. RPE1 PCM1 KO cells ciliated only up to 5.3%±0.9 compared to the 87.7% ±0.7 ciliating population in control cells after 48 h serum starvation, demonstrating an essential function for satellites in cilium assembly (Fig. S2A, B). This result is in agreement with the recent knockout study for PCM1 function in ciliogenesis (Wang et al, 2016). Unexpectedly, in contrast to RPE1 cells, lack of satellites resulted in a much smaller but still significant decrease in the fraction of ciliated cells in serum-starved IMCD3 cells (clone 1:55.5%±0.35, n=300, clone 2: 44.4%±3.3, clone 3:47.4%±0.02) relative to control cells (79.7%±2.46, n=300) (Fig. 2A, B). All three IMCD3 PCM1 KO clones had similar ciliogenesis defects (Fig. 2B) and we used clone # 1 for subsequent rescue and phenotypic characterization experiments (hereafter IMCD3 PCM1 KO). The ciliogenesis phenotype in IMCD3 PCM1 KO cells was rescued by stable expression of LAP-tagged full-length human PCM1, confirming that the ciliogenesis phenotype is a specific consequence of satellite loss (Fig. 2A, B). Despite the reduction in the ciliating population in IMCD3 PCM1 KO cells, the cilia that formed in these cells were normal in length (wt: 3.14µm±0.8, KO: 3.38µm±1.18), which suggests that PCM1 acts during cilia initiation (Fig. 2C).

**Figure 2.**
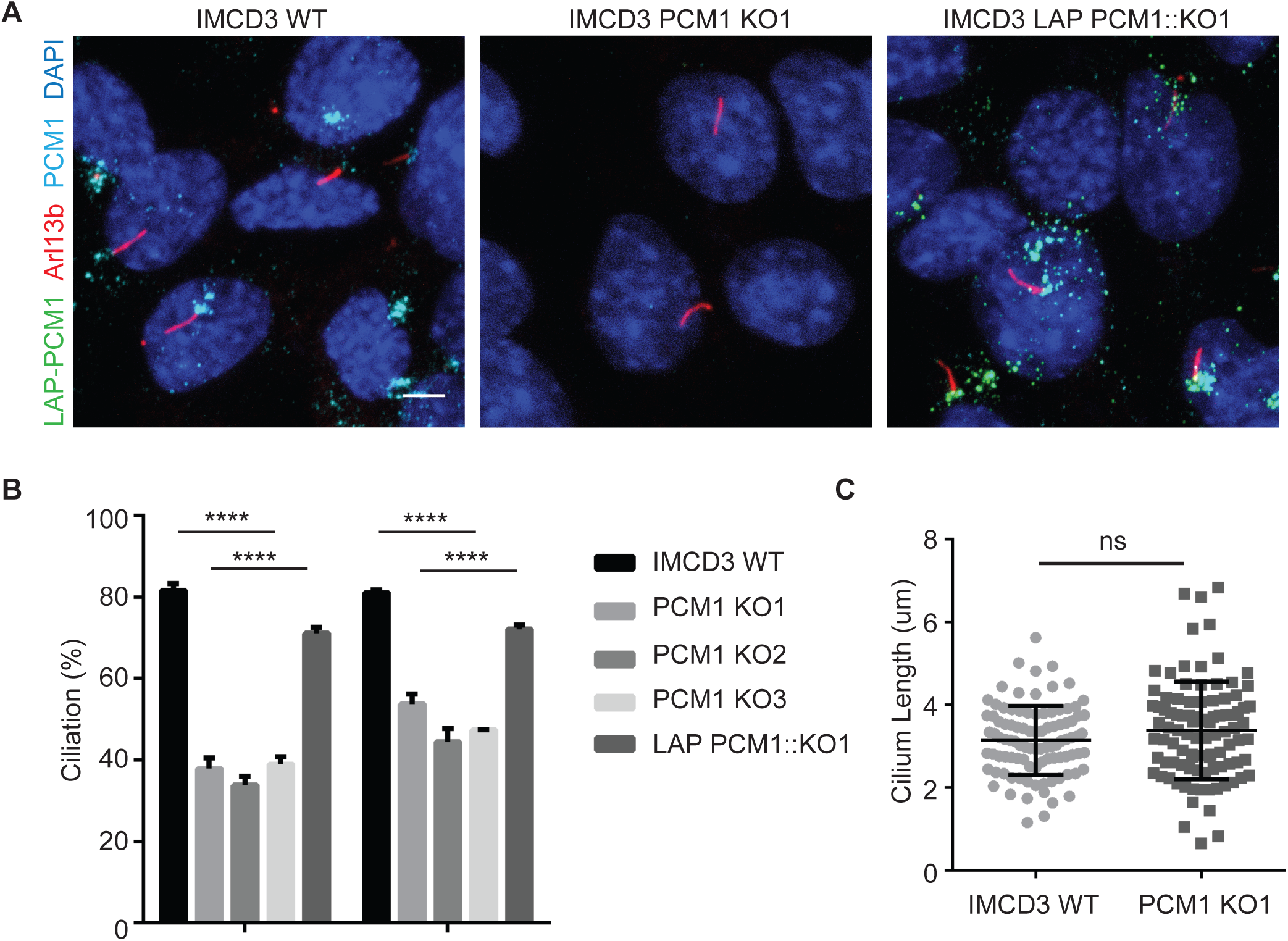
Satellites are required for efficient ciliogenesis. **A)** Effect of satellite loss on cilium formation. Control cells, IMCD3 PCM1 KO cells and LAP-PCM1-expressing IMCD3 PCM1 KO cells serum-starved for the indicated times and percentage of ciliated cells was determined by staining for acetylated-tubulin, Arl13b and DAPI. **(B)** Quantification of ciliogenesis and rescue experiments. Results shown are the mean of three independent experiments±SD (500 cells/experiment, ****p<0.0001, t test). **(C)** Effect of satellite loss on cilium length. IMCD3 cells serum-starved for 24 h and cilium length was determined by staining for acetylated-tubulin and DAPI.

We next asked whether the ciliogenesis defects of IMCD3 PCM1 KO cells are an indirect consequence of defects in cell cycle progression or centriole duplication. We tested the cell cycle-related phenotypes using a combination of assays. First, we performed proliferation assays with control and IMCD3 PCM1 KO cells by plating equal number of cells and counting them at indicated time intervals, which showed that loss of satellites did not have a significant impact on cell doubling times (Fig. 3A). Second, we analysed the fraction of control and IMCD3 PCM1 KO cells in different phases of the cell cycle using flow cytometry and showed that both had similar cell cycle profiles (Fig. 3B). Moreover, we performed live imaging of control and IMCD3 PCM1 KO cells stably expressing mCherry-H2B to assay for cell cycle progression defects. Both control and PCM1 KO cells had similar mitotic times (WT: 29 ±7.7min, KO: 28.1 ± 6.7min) (Fig. 3C, D). Finally, we quantified centrosome number in asynchronous control and IMCD3 PCM1 KO cells by gamma tubulin labelling and showed that PCM1 KO cells had similar centrosome counts relative to control cells (Fig. 3E). Collectively, these data indicate that satellites are not required for cell cycle progression and centriole duplication, and that the reduced ciliogenesis phenotype of PCM1 KO cells is not caused by defects in these processes.

**Figure 3.**
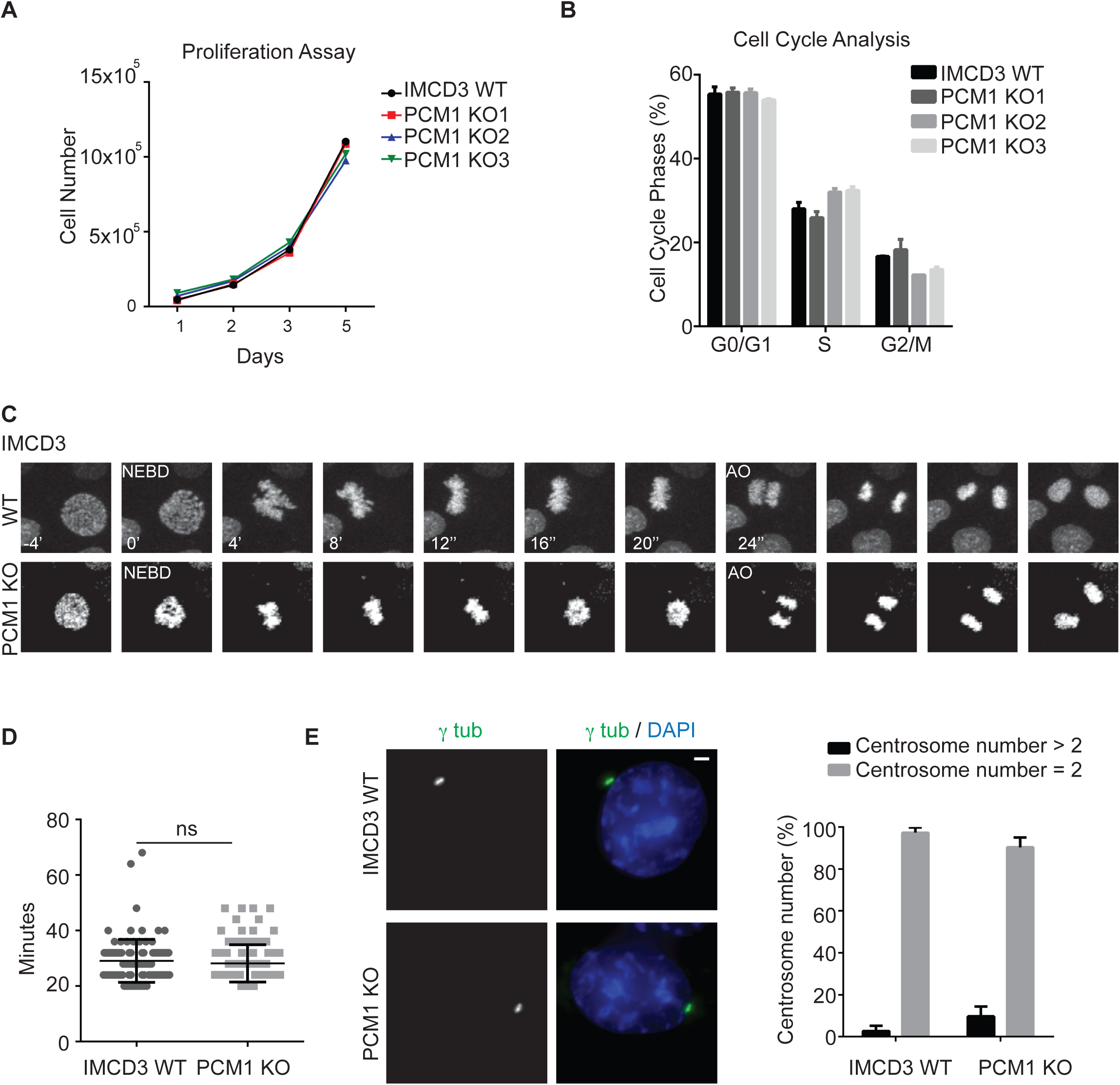
Satellites are not required for cell proliferation, cell cycle progression and centriole duplication. **(A)** Effect of satellite loss on percentage of cells in different cell cycle phases. Flow cytometry analysis of control or IMCD3 PCM1 KO cells. Data points show mean±SD of three independent experiments. **(B)** Effects of satellite loss on cell proliferation. 2*10^5^ cells were plated and counted at 24,48, 72 and 96 h. Data points show mean±SD of three independent experiments. There was no significant difference between control and IMCD3 PCM1 KO cells at any time point. **(C)** Effect of satellites loss on mitotic times. Control and IMCD3 PCM1 KO cell stably expressing mCherry-H2B were imaged every 6 min for 24 h. Mitotic time was quantified as the time interval from nuclear envelope breakdown (NEBD) to anaphase onset. Data points show mean±SD of two independent experiments. Control and IMCD3 PCM1 KO cells had similar mitotic times. **(D)** Representative still frames from time-lapse experiments were shown. **(E)** Effect of satellite loss on centriole duplication. Asynchronous control and IMCD3 PCM1 KO cells were stained with staining for gamma-tubulin and DAPI. Quantification results shown are the mean of two independent experiments±SD (100 cells/experiment, t test).

### Satellites have variable effects on regulating centrosomal and cellular abundance of key ciliogenesis factors

Satellites regulate centrosomal recruitment of proteins including centrin, pericentrin, ninein among others (Dammermann & Merdes, 2002). We hypothesized that the ciliogenesis defects of IMCD3 cells might be due to inefficient targeting of key ciliogenesis factors to the centrosome. To test this, we used quantitative immunofluorescence to determine the centrosomal levels of proteins that are components of distal appendages, transition zone proteins, IFT machinery as well as activators and suppressors of ciliogenesis in control and IMCD3 PCM1 KO cells.

Distal appendages function in docking of the mother centriole to the membrane during ciliogenesis and transition zone regulates the entry and exit of ciliary cargo (Mirvis et al, 2018; Wang & Dynlacht, 2018). The defects in ciliogenesis were not because of mislocalization of distal appendage or the transition zone proteins since Ceo164 and Cep290 localized correctly to the centrosomes in both control and IMCD3 PCM1 KO cells (Fig. 4A, B). Of note, the centrosomal levels of Cep164 was higher in PCM1 KO cells, which might be due to the increased abundance of Cep164 in total cell lysates in PCM1 KO cells (Fig. 4G). CP110 localizes to the distal ends of centrioles and is removed from the mother centrioles during ciliogenesis (Schmidt et al, 2009). In cycling cells, the centrosomal level of CP110 was similar in control and IMCD3 PCM1 KO cells (Fig. 4C). Importantly, IMCD3 PCM1 KO cells had a significant reduction in the basal body levels of IFT-B protein Ift88 (Fig. 4D). Given that the IFT-B machinery is required for the assembly of the ciliary axoneme (Bhogaraju et al, 2013; Haycraft et al, 2007; Pazour et al, 2002; Pazour et al, 2000), satellites might promote cilium assembly at a step upstream to the recruitment of the IFT machinery at the base of cilia.

**Figure 4.**
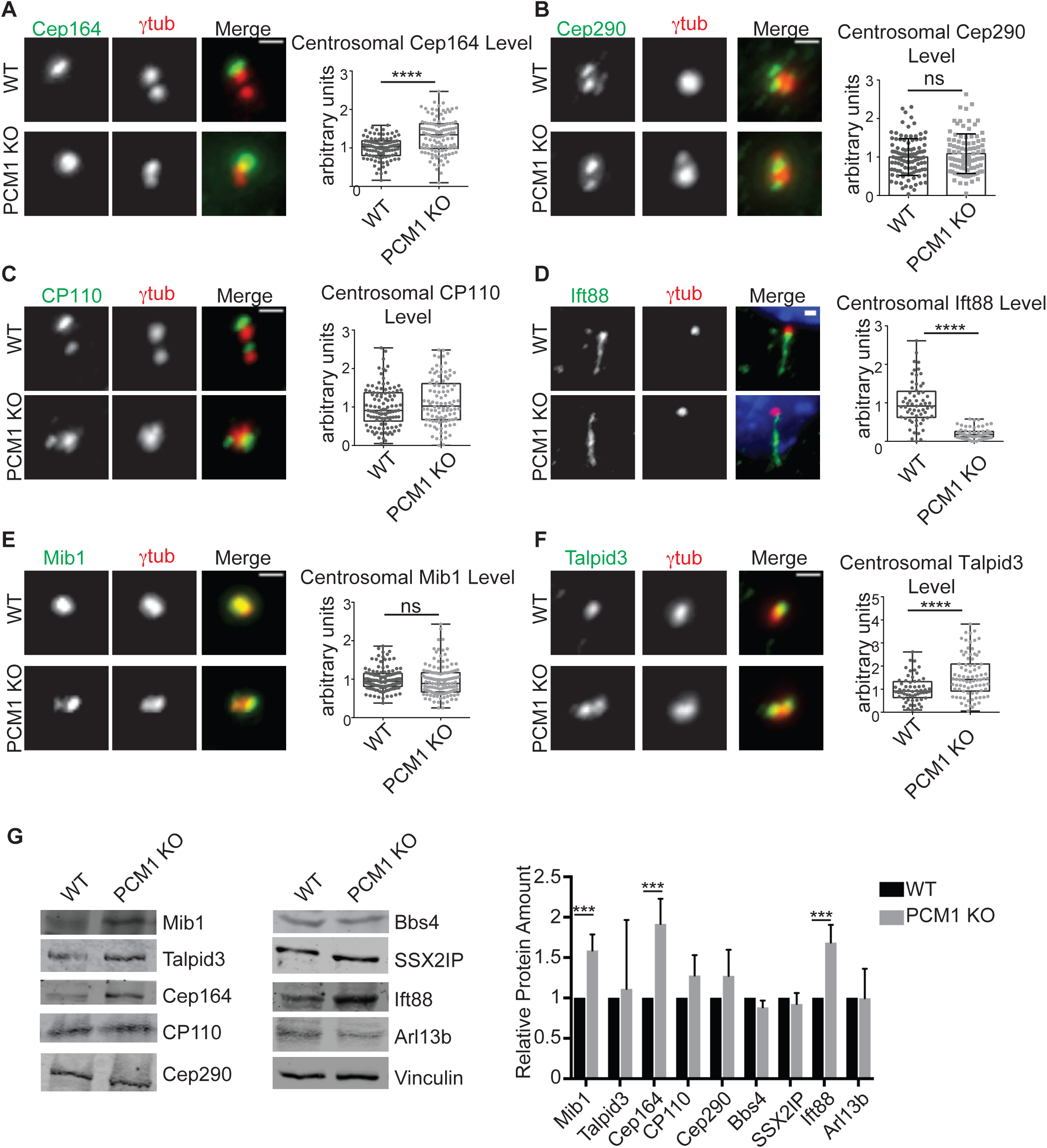
Satellites have varying degrees of effects on regulating centrosomal and cellular abundance of key ciliogenesis factors. **(A,B,C,D,E,F)** Effect of satellite loss on centrosomal abundance of proteins. Control and IMCD3 KO cells were fixed and stained with antibodies for the indicated proteins along with anti-gamma-tubulin and DAPI. Scale bar, 1 μm. Images represent cells from the same coverslip taken with the same camera settings. The centrosomal fluorescence intensity of the indicated proteins was measured for control and IMCD3 PCM1 KO cells in a 3 μm^2^ square area around the centrosome and levels are normalized to 1. Results shown are the mean of three independent experiments±SD (100 cells/experiment, **p<0.01, ***p<0.001, ****p<0.0001, ns: not significant, t-test)**(G)** Effects of satellite loss on cellular abundance of centrosome proteins. Whole-cell lysates from control and IMCD3 PCM1 KO cells were immunoblotted with the indicated antibodies. Vinculin was used as a loading control.

The inhibition of ciliogenesis in RPE1 PCM1 KO cells was shown to be a consequence of increased cellular and centrosomal levels of the E3 ubiquitin ligase Mib1, which led to a reduction in the cenrosomal levels of the key ciliogenesis factor Talpid3/KIAA0586 through its destabilization (Wang et al, 2016). We investigated whether loss of satellites had similar molecular consequences in IMCD3 cells. Despite the significant 1.5±0.26-fold increase in the cellular abundance of Mib1 in IMCD3 PCM1 KO cells (Fig. 4G), its centrosomal levels did not change in these cells relative to control cells (Fig. 4E). The centrosomal and cellular levels of Talpid3 in IMCD3 PCM1 KO cells were higher than control cells (Fig. 4F, G). Thus, in contrast to RPE1 cells, Mib1 sequestration and consequent Talpid3 degradation at the centrosome was not induced in satellite-less IMCD3 cells despite their increased cellular levels. Collectively, these results identify differences in the efficiency of the centrosomal targeting of key ciliogenesis factors as a likely mechanism for the phenotypic variability of satellite-less IMCD3 and RPE1 cells during cilium formation.

We also performed immunoblot analysis of whole-cell lysates of control and IMCD3 PCM1 KO cells to determine whether the cellular abundance of these proteins was affected. While there was an overall increase in the abundance of Mib1, Cep164 and Ift88, the abundance of Bbs4, SSXIIP, Cep290, Cp110, Talpid3 and Arl13b was unaffected (Fig. 4G). Thus, loss of satellites has differential effects on the abundance of different centrosome and cilium proteins, which was also the phenotype RPE1 PCM1 KO clones (Fig. S3C). Notably, except for Cep164, changes in abundance of these proteins were not reflected by similar changes in the recruitment of these proteins to the centrosome. For example, while Ift88 cellular levels increase significantly, its centrosomal levels had a significant reduction.

### Satellites regulate ciliary protein content

Analogous to their function in regulating protein targeting to the centrosome, we hypothesized that satellites might also regulate ciliary recruitment of proteins. To test this, we examined the composition of the ciliary shaft and the ciliary membrane of control and IMCD3 PCM1 KO cells by determining the ciliary levels of various proteins using quantitative immunofluorescence. Control and IMCD3 PCM1 KO cells had similar levels of ciliary Arl13b, which associates with the ciliary membrane via palmitoylation (Cevik et al, 2010) (Fig. 5A). However, IMCD3 PCM1 KO cells had significantly reduced ciliary levels of the IFT-B protein Ift88, somatostatin receptor 3 (Sstr3-LAP) and the serotonin 6 receptor (Htr6-LAP) (Fig. 5B, C, D). This reduction could be due to reduced diffusion between the cilia and the plasma membrane. To test this, we used half and full cilium fluorescence recovery after photobleaching (FRAP) (Hu et al, 2010) to measure the dynamic behaviour of HTR6-GFP in IMCD3 cells. However, the half time and percentage of recovery for HTR6-GFP was similar between control and KO cells (data not shown).

**Figure 5.**
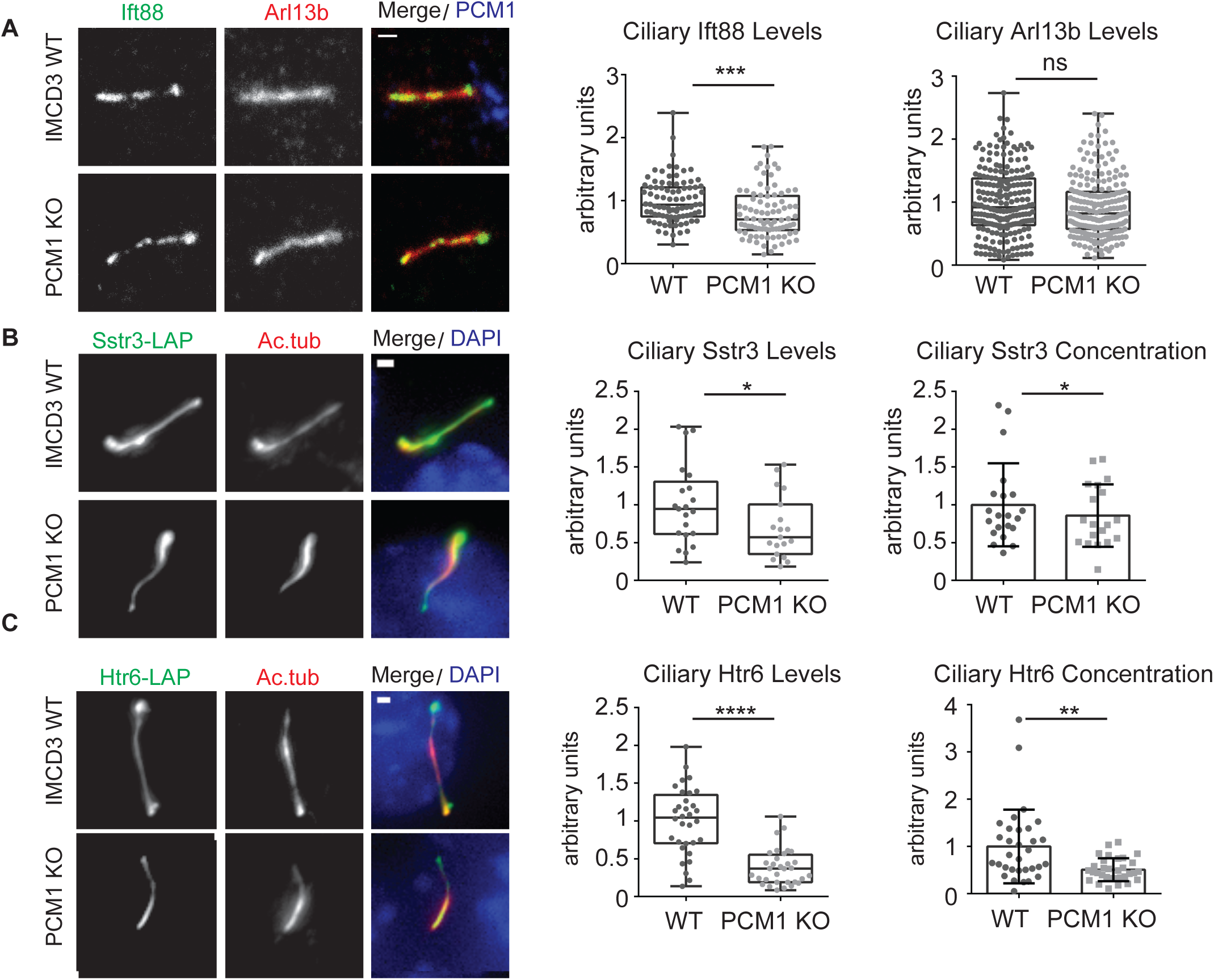
Satellites are required for regulating the ciliary content. Effects of satellite loss on the ciliary recruitment of ciliary membrane and shaft proteins. Control and IMCD3 PCM1 KO cells were serum starved for 24 h, fixed and stained for the cilium with anti-acetylated tubulin (Ac. tub) or Arl13b, and **(A)** Ift88 with anti-Ift88 and Arl13b with anti-Arl13b **(B)** Sstr3-LAP with anti-GFP, **(C)** 5Htr6-LAP with anti-GFP. **DNA was stained with DAPI.** Scale bar, 1 μm. Images represent cells from the same coverslip taken with the same camera settings. Relative quantifications of the ciliary intensity and ciliary concentration (fluorescence intensity/micrometer) of the indicated proteins were shown as the mean of two independent experiments±SD (100 cells/experiment, **p<0.01, ***p<0.001, ****p<0.0001, ns: not significant, t-test).

The ciliary localization of some Hedgehog pathway components is dynamically regulated in response to pathway activation. To elucidate the function of satellites in this dynamic regulation and to investigate whether the cilia assembled in IMCD3 satellite-less cells can still respond to Hedgehog ligands, we quantified the ciliary recruitment of the seven-transmembrane protein Smo, which moves to cilia in response to Sonic Hedgehog (Shh) ligands (Corbit et al, 2005). To this end, we stimulated control and IMCD3 PCM1 KO cells with 200 nM Smoothened agonist (SAG) in a time course manner and quantified the percentage of cilia with Smo. A significant (p<0.01) two-fold reduction in the percentage of cilia with Smo was observed in PCM1 KO cells (40.1%±7.9 of total) relative to control cells (80.6%±6.9 of total) at 4h, and 45.9%±18.2 of total in PCM1 KO cells relative to 75.5%±12.5 of total in control cells at 8h (Fig. 6A, B). Notably, control and IMCD3 PCM1 KO cells had similar percentages of cilia with Smo after 12 h and 24 h SAG stimulation, indicating that lack of satellites causes a delay in translocation of Smo to the cilium in response to SAG (Fig. 6A). We also analysed Smo levels at the cilia and found reduced localization in IMCD3 PCM1 KO cells (0.21%±0.15 of total) relative to control cells (1%±0.48 of total) after 4 h SAG treatment (Fig. 6B). This reduction was not due to changes in the cellular abundance of Smo (Fig 6C). To assay the transcriptional response to Hedgehog pathway activation, we examined the expression of the Hedgehog target gene *Gli1* in control and PCM1 KO cells after 24 h SAG stimulation by quantitative PCR. While wild-type cells had robust activation of Gli1 expression (normalized to 100%), PCM1 KO cells failed to upregulate Gli1 expression at 24 h (35%±25.6) (Fig. 6D). Taken together, these results indicate that satellites are required for the localization of sufficient levels of Smo at cilia, and efficient activation of the Hedgehog pathway.

**Figure 6.**
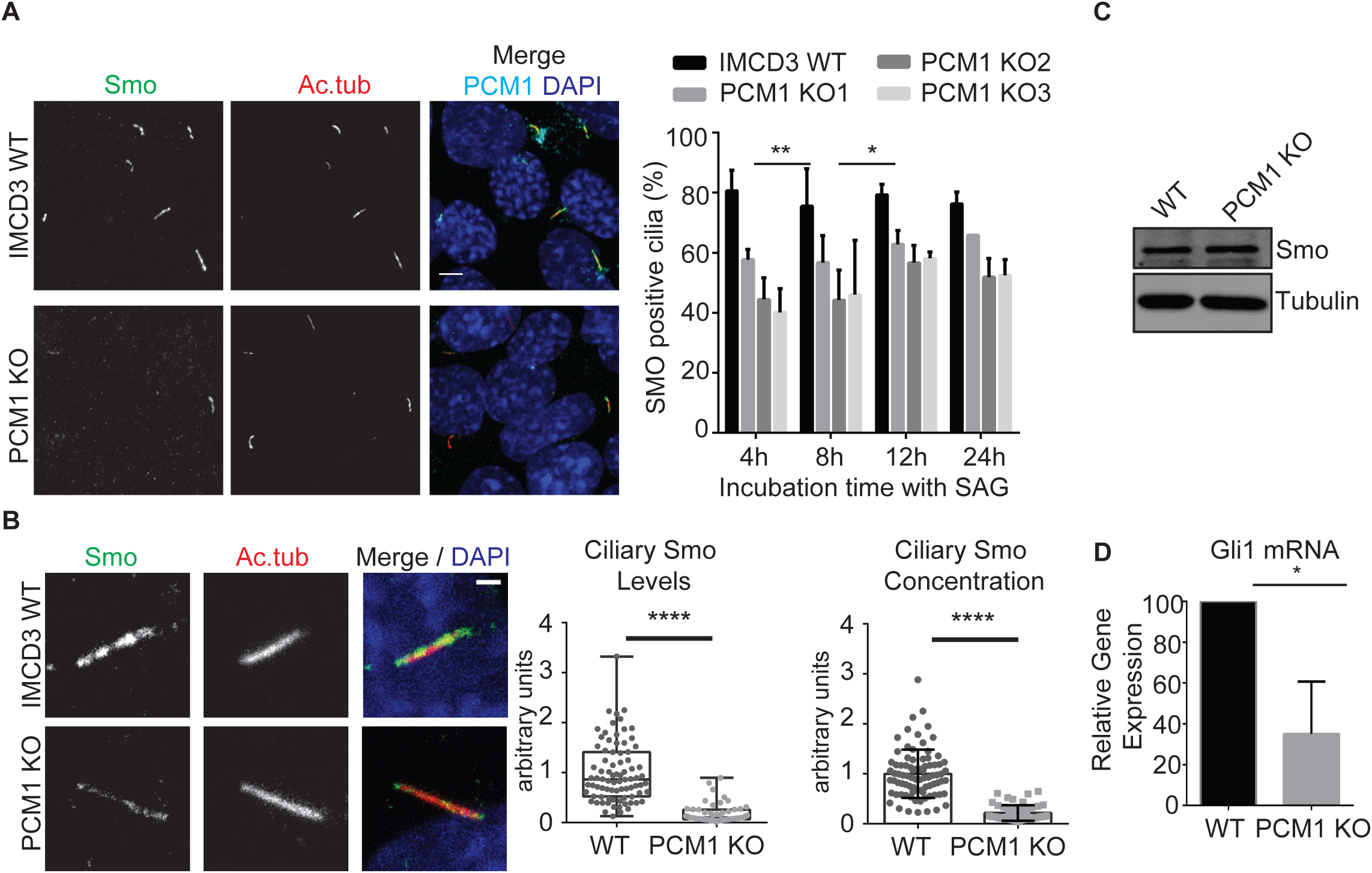
Satellites are required for ciliary Smo recruitment and Gli1 transcriptional activation in response to Hedgehog signals. **(A)** Effect of satellite loss on ciliary recruitment of Smo. Control and IMCD3 PCM1 KO cells were serum starved for 24 h, treated with 200 nM SAG for the indicated times, fixed and stained for Smo, acetylated tubulin (Ac. tub) and DAPI. Percentage of Smo-positive cilia was quantified. Scale bar, 4 μm. Results shown are the mean of three independent experiments±SD (250 cells/experiment, **p<0.01 *p<0.5, t test). **(B)** Effect of satellite loss on ciliary SMO levels. The ciliary intensity of SMO was quantified by staining 4 h SAG-treated cells for Smo and acetylated tubulin. DNA was stained with DAPI. Relative quantifications of the ciliary intensity and ciliary concentration of SMO was shown as the mean of two independent experiments±SD (100 cells/experiment, ****p<0.0001 t-test). **(C)** Effect of satellite loss on cellular abundance of Smo. Whole-cell lysates from control and IMCD3 PCM1 KO cells treated with SAG for 24 h were immunoblotted with Smo and alpha tubulin (loading control). Smo cellular levels were similar between control and IMCD3 PCM1 KO cells. **(D)** Effect of satellite loss on Gli1 transcriptonal activation. Gli2 mRNA was quantified by qPCR and its fold change is normalized to control cells (=100). Results shown are the mean of two independent experiments±SD (*p<0.05, t-test).

### Satellites are required for epithelial cell organization in 3D cultures

When grown in a three-dimensional gel matrix, epithelial cells organize into polarized, spheroid structures that reflect the *in vivo* organization of the epithelial tissues. Epithelial spheroids have been widely used to assay cilia dysfunction, because proper cilium assembly and ciliary signalling is essential for the establishment of the highly organized architecture and apicobasal polarity of epithelial cells in 3D (Delous et al, 2009; Mahjoub & Stearns, 2012; Otto et al, 2010). To assay the consequences of ciliary defects associated with loss of satellites on tissue architecture, we used the 3D spheroid cultures of IMCD3 cells that mimics *in vivo* organization of the kidney collecting duct (Giles et al, 2014). Control and IMCD3 PCM1 KO cells were grown in Matrigel for 3 days, serum starved for 2 days and spheroid architecture was visualized by staining cells for markers for cilia, cell-cell contacts and polarity. Control cells formed spheroids with prominent lumen, apical cilia oriented towards the lumen, and proper organization of apical and basal surfaces (ZO-1 and beta-catenin) (Fig. 7A). However, PCM1 KO cells had a significant decrease in their ability to form proper spheroids relative to control cells (64.9%±2.97 spheroids in control cells compared with 40.87%± 9.42 for PCM1 KO cells; p=0.0135) (Fig. 7B). Instead, they formed cell clusters with no lumens that had misoriented cilia and disorganized apical and basolateral junctions (Fig. 7A). In agreement with the function of satellites in ciliogenesis and signalling, these results identify important roles for satellites in epithelial organization in this *in vitro* tissue model.

**Figure 7.**
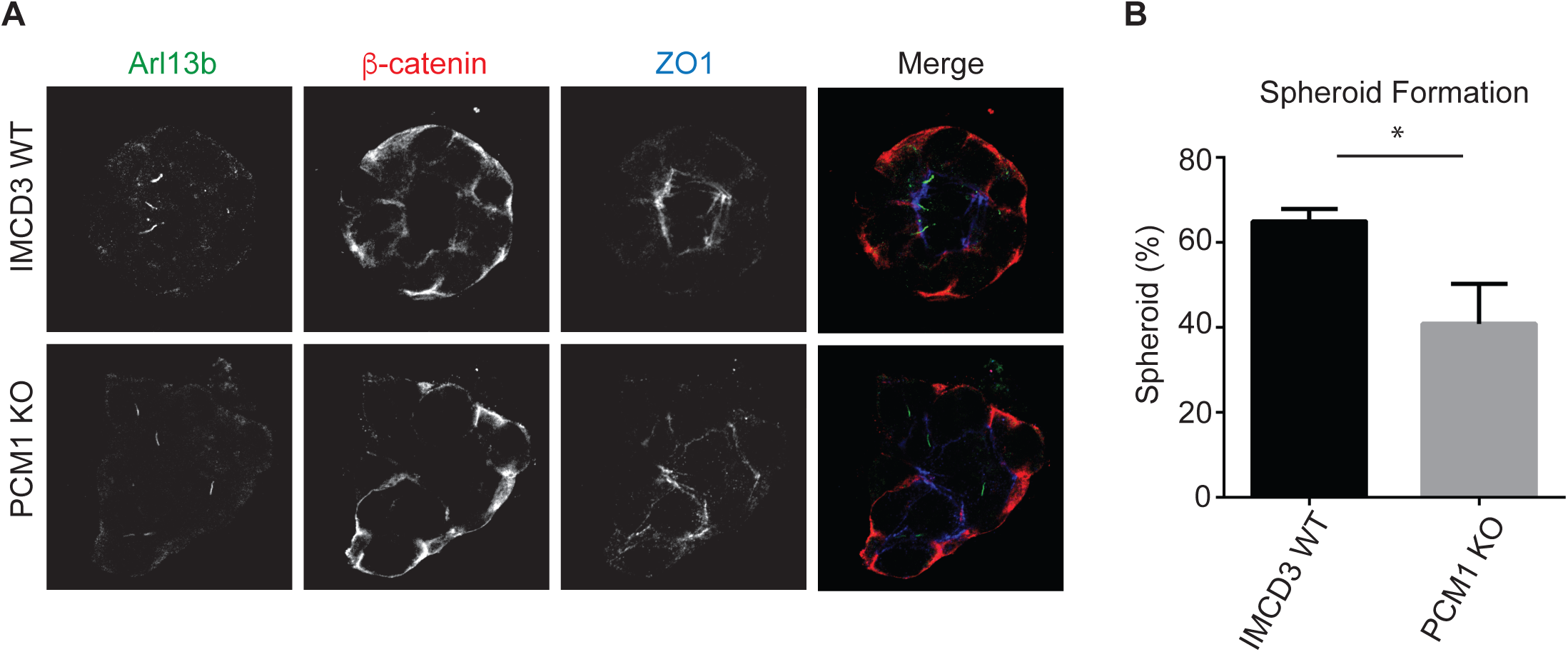
Satellites are required for epithelial cell organization in 3D spheroid cultures. **(A)** Effects of satellite loss on epithelial cell organization. IMCD3 cells were grown in Matrigel for 3 days and serum starved for 2 days. The spheroids were fixed and stained for apical junctions with anti-ZO1, cilia with anti-Arl13b and basolateral surfaces with anti-beta-catenin. Scale bar, 10 μm. **(B)** Quantification of the frequency of spheroid formation in control and IMCD3 PCM1 KO cells. Results shown are the mean of three independent experiments±SD (50 spheroids/experiment, *p<0.05, t-test).

### Loss of satellites leads to a rearrangement of the global proteome profile, but not the global transcriptome profile

Loss of satellites variably affected the cellular abundance of centrosome and cilium proteins both in IMCD3 cells and RPE1 cells (Fig. 4, S2) (Wang et al, 2016). This suggests functions for satellites in regulating the cellular abundance of proteins. However, whether these changes are a consequence of transcriptomic or proteomic modulation and which centrosome proteins are affected by these changes have not been investigated. To address this, we performed a systematic comparative analysis of the global transcriptomic and proteomic profile of IMCD3 and RPE1 PCM1 KO cells.

RNA sequencing (RNAseq) analysis of PCM1 KO and control RPE1 and IMCD3 cells revealed a small number of genes, 8 in IMCD3 PCM1 KO cells and 23 in RPE1 PCM1 KO cells, which were differentially regulated with log_2_ fold change >1.5 and p-value <0.05 (Table 1, S1). Gene ontology and KEGG pathway functional analysis did not identify any functional clustering among these genes. Moreover, none of the differentially regulated genes were shared between IMCD3 and RPE1 cells. Therefore, it seems unlikely that the differential expression of the identified genes is a direct result of the loss of satellites and the direct cause of the phenotypes observed in PCM1 KO cells. Together, these results show that loss of satellites does not induce a major stress response or compensation mechanism in these cells.

**Table 1.** Differentially regulated genes in IMCD3 PCM1 KO cells compared to control cells (>1.5-fold, adj.P-value < 0.05) Genes significantly up- or down-regulated in RPE1 PCM1 KO cells compared to WT controls (>1.5-fol, adj. P-value<0.05)

| Gene Symbol | Gene Name | Function | Regulation | Fold Change |
| --- | --- | --- | --- | --- |
| Pkrd1 | protein kinase D1 | Angiogenesis, Apoptosis, Differentiation | DR | Inf |
| Tsnax | translin-associated factor X | Differentiation, Spermatogenesis | DR | Inf |
| B4galt4 | UDP-Gal:betaGlcNAc beta 1,4-galactosyltransferase, polypeptide 4 | Protein glycosylation | DR | 15.2 |
| Fstl1 | follistatin-like 1 | Response to starvation | DR | 12.7 |
| Fat4 | FAT atypical cadherin 4 | Cell adhesion | DR | 10.2 |
| Mlf1 | myeloid leukemia factor 1 | Cell cycle, Differentiation | DR | 9.4 |
| Dmd | dystrophin, muscular dystrophy | Muscle cell cellular homeostasis | DR | 6.1 |
| Pcm1 | pericentriolar material 1 | Cilium biogenesis | DR | 4.3 |
| Gxylt2 | glucoside xylosyltransferase 2 | O-glycan processing | DR | 3.7 |

In contrast to the transcriptome analysis, tandem mass tag (TMT) labelling-based quantitative analysis of the global proteome of control and IMCD3 PCM1 KO cells showed that loss of satellites substantially altered the proteomics profile. A total of 6956 proteins were identified with high confidence, 295 of which were upregulated, while 296 were downregulated (log_2_-transformed normalized ratios>0.5) (Fig. 8A). GO-term analysis of the major biological processes and compartments enriched or depleted significantly in PCM1 KO cells identified categories related to “centrosome”, “cell proliferation” and “microtubules” (Fisher’s exact test, FDR<0.05) (Fig. 8B, S3). This is in agreement with the function of satellites as regulators of the centrosome/cilium-complex. Importantly, the GO terms actin cytoskeleton, cell migration and adhesion, endocytosis, neuronal processes, metabolic processes were among the most enriched and depleted compartments and processes (Fig. 8B), which was also illustrated as functional network clustering analysis in Fig. S3. Together, these analysis suggest possible functions for satellites in these processes (Fig. 8B)

**Figure 8.**
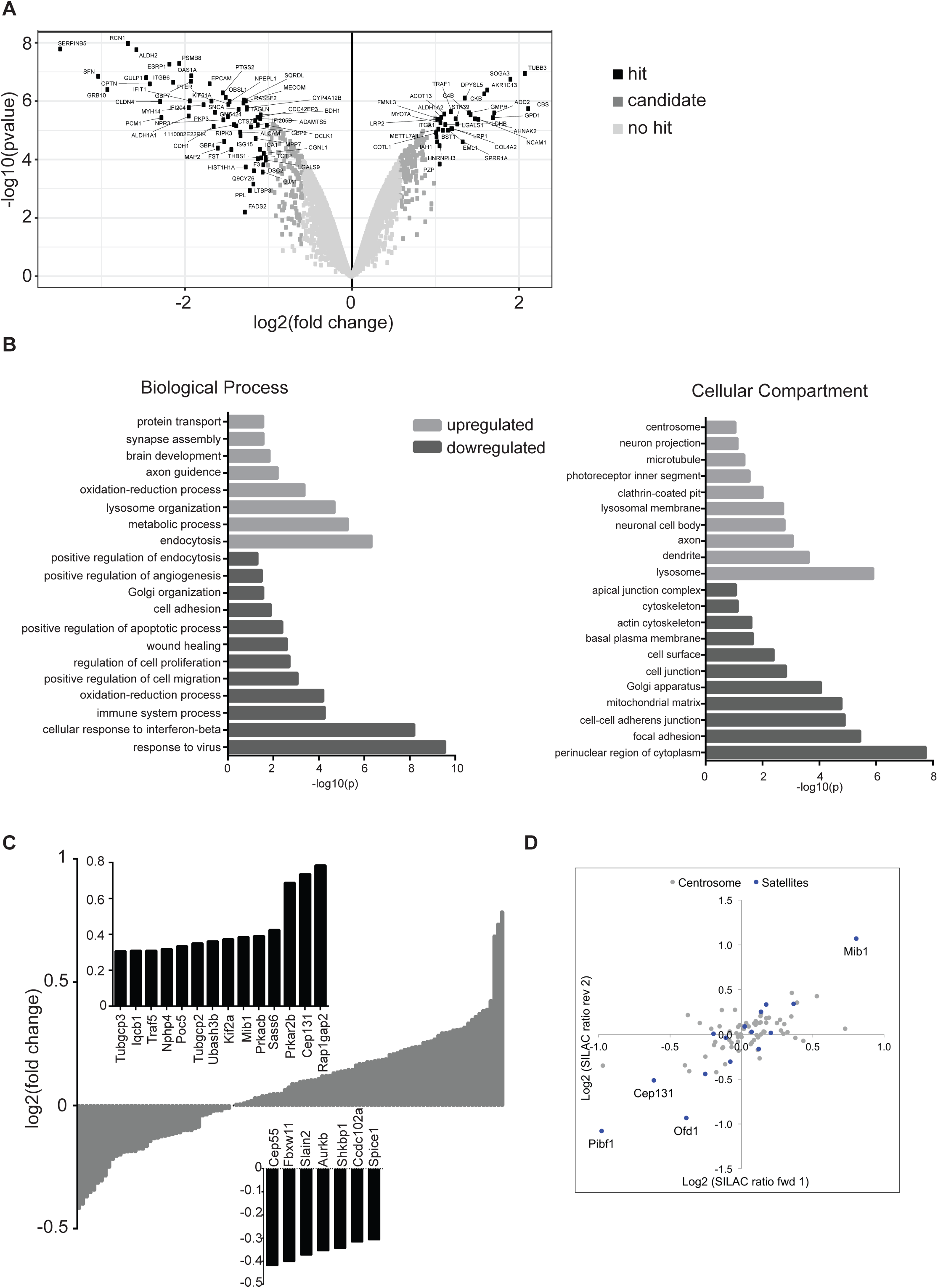
Loss of satellites changes the global proteome. Quantitative proteomics profiling of IMCD3 and RPE1 PCM1 KO cells. **(A)** Volcano plots illustrating that 591 proteins show a significant change in abundance. Data was derived from mass spectrometry analysis of three biological replicates of control and IMCD3 PCM1 KO cells. The x-axis corresponds to the log_2_ fold change value, and the y-axis displays the −log_10_ P value. Proteins were classified into hits (indicated with black circles) if the FDR is smaller than 0.05 and a fold change of at least 2-fold is observed. Proteins were classified into candidates (indicated with medium gray circles) if the FDR is smaller than 0.2 and a fold change of at least 1.5 fold is observed. **(B)** GO-term enrichment analysis of upregulated and downregulated proteins based on biological process and cellular compartment **(C)** Comparison of the centrosome proteome with the IMCD3 PCM1 KO global proteome. 112 centrosome proteins identified in the global proteome (x-axis) was plotted against their log_2_-transformed ratios of fold changes (y-axis) between IMCD3 PCM1 KO and control samples. The insets show magnified views of the significantly enriched and depleted proteins (log_2_>0.3 and log_2_<-0.3). **(D)** Abundance changes of centrosome proteins identified in the SILAC global proteome of RPE1 PCM1 KO and control cells. The fold change in SILAC protein ratios for the centrosome proteins (gray circles) identified in the global proteome were plotted for forward and reverse SILAC experiments. Satellite proteins that are significantly altered in expression (log_2_>0.5 and log_2_<-0.5) were indicated as blue circles.

To specifically determine the consequences of satellite loss on the cellular abundance of centrosome proteins, we cross-correlated the global proteome dataset with the published centrosome proteome (Jakobsen et al, 2011). Unexpectedly, we found that majority of the 116 centrosome proteome identified remains in place, where only a few proteins including the satellite protein Cep131 and regulatory proteins Ran1Gap2 and Prkar2B were perturbed in the global proteome (Fig. 8C) (Sadrian et al, 2012; Sha et al, 2018; Sha et al, 2017; Staples et al, 2012). To investigate whether RPE1 PCM1 KO cells had similar changes, we determined the global proteome of these cells using SILAC-based mass spectrometry analysis. Like IMCD3 cells, in RPE1 PCM1 KO cells, only a few of the centrosome proteins including all satellite proteins Mib1, Ofd1, Cep131 and Pibf1/Cep90 were substantially altered in expression while the remaining 77 centrosome proteins were unaffected (log_2_>0.5 for SILAC labelling) (Fig. 8D). Due to low abundant nature of centrosome proteins, we set a lower significance threshold (log_2_>0.3 for TMT labelling) for detecting changes in abundance, which identified 21 proteins in IMCD3 KO cells to be differentially expressed. The upregulated and downregulated centrosome proteins were implicated in a wide range of processes including suppressors and activators of ciliogenesis, centriole duplication, spindle formation, signalling and microtubule cytosketolon (Fig. 8C). The lack of functional clustering among these proteins shows that the regulatory roles of satellites on their resident proteins are specific to each protein and that phenotypic consequences of loss of satellites are likely combinatorial effects of these changes, instead of a specific pathway.

## Discussion

The results of our study reveal three specific roles of satellites in vertebrate cells. First, satellites are required for efficient ciliogenesis, and this requirement varies across different cell types. Second, satellites regulate the composition of the ciliary shaft and the ciliary membrane and these functions are required for epithelial cell organization in 3D cultures. Finally, satellites are regulators of the cellular proteostatis of not only centrosome proteins but also proteins from a diverse set of pathways. Given that mutations affecting various satellite components cause ciliopathies (Lopes et al, 2011; Tollenaere et al, 2015) and loss of satellites in zebrafish leads to ciliopathy-related phenotypes (Stowe et al, 2012), our findings suggest variable degrees of defects in cilium assembly, signalling and tissue architecture linked to satellite mutations as mechanisms underlying diverse disease phenotypes of ciliopathies, in particular in retinal degeneration and polycystic kidney disease.

A variety of satellite-specific functions were reported in previous studies through loss of function studies targeting PCM1. Transient depletion of PCM1 caused defects in organization of the interphase microtubule network in U2OS cells (Dammermann & Merdes, 2002), cell cycle progression and cell proliferation in MRC-5 primary human fibroblasts and HeLa cells (Srsen et al, 2006), and cilium formation in RPE1 cells (Kim et al, 2008; Nachury et al, 2007). Constitutive depletion of PCM1 from RPE1 and GBM cells caused inhibition of ciliogenesis (Hoang-Minh et al, 2016; Wang et al, 2016). We found that satellites are only required for cilium assembly and function, but not for cell proliferation, cell cycle progression and centriole duplication in IMCD3 cells. This variation in phenotypes, except for the cilium-related ones, could be due to a functional compensation mechanism that rescues the defects caused by constitutive loss of satellites, previously reported for another satellite protein Azi1 (Hall et al, 2013). Alternatively, the requirement for satellites in these processes may simply vary in different cell types, as we show for cilium-related processes in this study. We would like to highlight that loss of function studies of PCM1 also defined functions for satellites in autophagy (Joachim et al, 2017), stress response (Nielsen et al, 2018; Villumsen et al, 2013), and neuronal progenitor cell proliferation and migration (Ge et al, 2010; Insolera et al, 2014; Zhang et al, 2016), which we have not focused on in this study.

Cell-type specific functions of satellites, in particular during cilium formation and function, have not previously been addressed. Satellite loss in RPE1 cells caused an almost complete inhibition of ciliogenesis, identifying a direct and requisite role for satellites in cilium assembly in these cells (Wang et al, 2016). In contrast, satellite-less IMCD3 cells had a much weaker two-fold reduction in their ability to ciliate. IMCD3 cells that ciliated in the absence of satellites had cilia of similar length compared to control cells, which suggests that satellites in these cells are required to initiate assembly of the ciliary axoneme. While satellite-less cells proceeded normally through early steps of mother centriole maturation, they were defective in the recruitment of Ift88 to the basal body, suggesting that satellites promote ciliogenesis upstream of this stage during ciliogenesis. Supporting the relationship between IFT machinery and satellites, proteomics studies identified putative interactions of satellites with various IFT-A and IFT-B components, supporting the functional relationship between the IFT machinery and satellites (Gupta et al, 2015; Huttlin et al, 2017; Huttlin et al, 2015). Of note, we observed that centrosomal levels of Cep164 and Talpid3 increased in satellite-less cells and we do not know whether these changes have any functional consequences on ciliogenesis.

The mechanisms underlying the cell-type specificity of satellite functions are unknown. Ciliogenesis pathways vary among different cell types, which might cause the differential requirements for satellites during cilium assembly. While most cells including RPE1 cells use the intracellular pathway, in which the basal body interacts with ciliary vesicles and begins axoneme elongation in the cytosol; polarized epithelial cells like IMCD3 cells use the extracellular pathway, in which basal body docks to the apical plasma membrane to induce axoneme elongation (Mirvis et al, 2018; Wang & Dynlacht, 2018). Given that there are major differences in the stages of ciliogenesis between IMCD3 and RPE1 cells, these cells might have differential molecular requirements for regulating these events. In agreement, we identified differences in how loss of satellites is reflected in the centrosomal and cellular levels of key ciliogenesis proteins including Mib1 and Talpid3, which likely contributes to the phenotypic variability of satellite-less retinal and kidney epithelial cells during cilium assembly.

Satellites were shown to regulate the stability of the ciliogenesis factor Talpid3 and the Atg8 autophagy marker Gabarap through sequestration of Mib1, which suggested that they might mediate functions as regulators of protein degradation pathways (Joachim et al, 2017; Wang et al, 2016). However, the turnover of which centrosome proteins are regulated by satellites was not known. Global proteomics of RPE1 and IMCD3 cells showed that the majority of the centrosome proteome remained unaltered and that only the expression of a small subset of centrosome proteins were perturbed. Importantly, Mib1 was among the most upregulated centrosome proteins in both RPE1 and IMCD3 satellite-less cells, corroborating the negative crosstalk between PCM1 and Mib1 stability (Akimov et al, 2011; Joachim et al, 2017; Villumsen et al, 2013; Wang et al, 2016). It is important to note that the global proteome analysis is limited in identifying all the centrosome proteins and determining changes in their expression levels with high sensitivity, due to the low abundance levels of most centrosome proteins in cells (Bauer et al, 2016).

In addition to regulating protein stability, satellites were shown to mediate their functions through targeting proteins to the centrosome or sequestering them to limit their centrosomal recruitment. In agreement, we detected changes in the centrosomal and ciliary levels of key ciligoenesis proteins, as well as ciliary membrane and shaft proteins. In addition to showing functions for PCM1 in the ciliary recruitment of proteins for the first time, these defects provide insight into the mechanisms underlying satellite-related defects *in vivo* and in disease Notably, for some proteins like Ift88 and Mib1, we showed that changes in the centrosomal levels of these proteins do not scale with changes in their cellular abundance. An attractive possibility for this discrepancy is that satellite-less cells compensate for functional defects through modulating the centrosomal recruitment of proteins. Because we performed this analysis only for a handful of proteins, future studies like identification of the centrosome and cilium proteome in satellite-less cells are required to elucidate the complete list of centrosome and cilium proteins regulated by PCM1.

Despite the unaltered abundance of the centrosome proteome, loss of satellites led to a significant rearrangement of the global proteome, suggesting that satellites regulate cellular proteostasis. Importantly, these global studies provided insight into previously uncharacterized satellite functions. Among the most enriched pathways were the ones related to the actin cytoskeleton such as cell migration, cell adhesion and endocytosis. The possible functional connection of satellites to actin-related processes is noteworthy due to the function of centrosomes as actin-nucleating centres. In particular, satellites are required for centrosomal actin nucleation through targeting actin assembly factors to the centrosome (Farina et al, 2016; Obino et al, 2016). Satellites might also function in other actin-regulated ciliary processes, such as periciliary vesicle transport to the basal body and cilia length regulation (Kim et al, 2015; Kim et al, 2010; Kohli et al, 2017; Wu et al, 2018). Moreover, we also found a markedly strong increase at the protein level in neurogenesis-linked pathways, such as neuronal body and extensions, postsynaptic membrane and axon guidance. Consistently, mutations in PCM1 were associated with schizophrenia (Datta et al, 2010; Gurling et al, 2006; Kamiya et al, 2008), and PCM1 and satellite proteins, Hook3, DISC1 and SDCCAG8 were implicated in maintaining neural progenitor cells and neuronal migration during cortical development. Future studies on dissecting the function of satellites in actin-related cellular processes and neurogenesis will be important to identify the full range of satellite functions.

While phenotypic characterization of stable genetic knockout model for of satellite-less cells have led to important insights into satellite function and molecular mechanism, its disadvantage is the possible masking of some satellite phenotypes. Generation and maintenance of PCM1-/-lines using the CRISPR/Cas9 genome editing approach requires a long time frame where cells undergo many successive divisions without satellites and thus might have enough time to activate compensatory mechanisms that can mask satellite-specific phenotypes, specifically through proteomic modulation for satellite-less cells as their transcriptome scarcely change (El-Brolosy & Stainier, 2017; Rossi et al, 2015). Likewise, some of the changes we detected in global transcriptomics and proteomics might be specific to cell clones. Future studies using spatially and temporally controlled *in vitro* and *in vivo* experiments are required to overcome these limitations.

In summary, our findings identify a more complex molecular and functional relationship between satellites and the centrosome/cilium complex. We suggest that satellites are among the additional regulatory mechanisms vertebrates acquired during evolution in order to prevent defects in the biogenesis and function of the centrosome/cilium complex and thereby associated diseases. Future studies are required to elucidate the functions of satellites in different cell types and tissues and to investigate how defects in these functions are associated with different disease states.

## Materials and Methods

### Cell Culture and Transfection

RPE1-hTERT and IMCD3::Flippin cells (gift from Max Nachury, UCSF, CA) were grown in DMEM/F12 50/50 (Life Technologies) supplemented with 10% fetal bovine serum (FBS; Life Technologies). HEK293T cells were grown in DMEM supplemented with 10% FBS. Media was supplemented with 100 U/ml penicillin and 100 µg/ml streptomycin (Life Technologies). All cells were cultured at 37 °C and 5% CO_2_. RPE1 and IMCD3 cells were transfected with the plasmids using Lipofectamine LTX according to the manufacturer’s instructions (Life Technologies). HEK293T cells were transfected with the plasmids using 1 μg/μl polyethylenimine, MW 25 kDa (PEI, Sigma-Aldrich, St. Louis, MO). IMCD3 cells stably expressing HTR6-LAP, SSTR3-LAP, LAP-PCM1 and LAP-PCM1 (1-3600) were generated by cotransfeting IMCD3 Flip-in cells with pEF5.FRT.LAP expression vectors and pOG44. For cilium-related experiments, cells were incubated in low-serum medium (0.5% serum) for 24 h or 48 h. For Smo relocalization experiments, cells were serum starved for 24 h and incubated with 200 nm SAG (EMB Milipore) for 4, 8, 12 or 24 h.

### 3D Spheroid Assay

40 ul/well of 100% Matrigel (BD Biosciences) were solidified in Lab-Tek 8-well chamber slides (Thermo Fisher) through 15 min incubation at 37º. Control and IMCD3 PCM1 KO cells were trypsinized, washed with PBS and resuspended in DMEM/F12 50/50 supplemented with 2% fetal bovine serum and 2% Matrigel and 5000 cell/well were plated in Matrigel-coated 8-well chamber slides. Cells were grown at 37º for 4 days, serum-starved for 1 day, fixed with paraformaldehyde (PFA) and stained with the indicated markers. Cell clusters with no hollow lumen, multiple lumens, disturbed organization of apical and basal markers and/or misaligned nuclei were quantified as defective.

### Lentivirus production and cell transduction

Lentivirus were generated as described (Mahjoub et al, 2010), using pLentiCRISPRv2, pLentiCRISPRv2-mousePCM1 and pLentiCRISPRv2-humanPCM1 plasmids as transfer vectors. For infection, 1×10^5^ RPE1 cells were seeded on 6-well tissue culture plates the day before infection, which were infected with 1 ml of viral supernatant the following day. 24 h post-infection, medium was replaced with complete medium. 48 hours post-infection, cells were split and selected in the presence of 6 µg/ml puromycin for RPE1s and 2 ug/ml puromycin for IMCD3 cells for five to seven days till all the control cells died. LentiCRISPRv2 empty vector-transfected and puromycin-selected cells were used as control. After the PCM1 knockout efficiency was determined in the heterogeneous pool using immunofluorescence experiments, cells were trypsinized and serial dilutions were performed into normal growth medium. Colonies formed within 10-14 days were then trypsinized and expanded for screening for knockouts by immunofluorescence and western blotting. IMCD3 cells stably expressing mCherry-H2B were generated by infection of cells with mCherry-H2B-expressing lentivirus. Lentivirus were generated using pLVPT2-mCherry-H2B plasmid as transfer vectors.

### Cloning and Genome editing

Full-length cDNA of BBS4 (NM_001252678) was obtained from DF/HCC DNA Resource Core (Harvard Medical School, MA). Full-length cDNAs for human PCM1 (NM_006197), HTR6 (NM_021358) and SSTR3 (BC120843) were amplified from the peGFP-N1-PCM1, peGFP-N1-SSTR3 and peGFP-N1-HTR6 plasmids. PCR-amlified open reading frames of full-length PCM1 and its deletion mutant (1-3600), HTR6, BBS4 and SSTR3 were cloned into pDONR221 using the Invitrogen Gateway system. Subsequent Gateway recombination reactions using pMN444 pEF5-FRT-LAP DEST, provided by M. Nachury (UCSF, CA), was used to generate LAP-PCM1, LAP-PCM1(1-1200), HTR6-LAP, SSTR3-LAP and BBS4-LAP. Guide RNAs targeting human and mouse PCM1 coding exon 2 were cloned into LentiCRISPRv2 (gift of Feng Zhang, Addgene plasmid #52961) using the following primers: mouseOligo forward: 5′-

CACCGCTGCTGTGTGGAAACGTATG -3′, mouseOligo reverse: 5’-

AAACCATACGTTTCCACACAGCAGC-3′, humanOligo forward: 5′-

CACCGCTACTGTGTGGGAACGTATG -3′, humanOligo reverse: 5’-

AAACCATACGTTCCCACACAGTAGC-3′, nontargeting forward:

5’-CGCTTCCGCGGCCCGTTCAA-3′ -3′, nontargeting reverse:

5’TTGAACGGGCCGCGGAAGCG-3′ -3′ All plasmids were verified by DNA sequencing.

### Flow cytometry and proliferation assays

Cells were harvested and fixed in 70% ethanol at -20 overnight, followed by washes in phosphate-buffered saline (PBS) and staining with 40 µg g/ml propodium iodide and 10 µg g/ml RNaseA at 37 for 30 min. Cell cycle analysis was performed using a Sony SH800S Cell Sorter (Sony). For proliferation assays, 2×10^5^ cells were plated and cells were counted at 24, 48, 72 and 96 h. At each count, cells were splitted at 1:2.

### Cell lysis and immunoblotting

For preparation of cell extracts, cells were lysed in 50 mM Tris (pH 7.6), 150 mM NaCI, 1% Triton X-100 and protease inhibitors for 30 min at 4°C and centrifuged at 15.000 g for 15 min. The protein concentration of the resulting supernatants was determined with the Bradford solution (Bio-Rad Laboratories, CA, USA). For immunoblotting, cell extracts were first resolved on SDS-PAGE gels, transferred onto nitrocellulose membranes and blotted with primary and secondary antibodies. Visualization of the blots was carried out with the LI-COR Odyssey^®^ Infrared Imaging System and software at 169 µm (LI-COR Biosciences).

### RNA isolation, cDNA Synthesis and qPCR

Total RNA was isolated from control and IMCD PCM1 KO cells after 24h SAG treatment using NucleoSpin RNA kit (Macherey-Nagel) according to the manufacturer’s protocol. Quantity and purity of RNA was determined by measuring the optical density at 260 and 280 nm. Single strand cDNA synthesis was carried out with 1 mg of total RNA using iProof High-Fidelity DNA Polymerase) qPCR analysis of Gli1 was performed with primers 5’ GCATGGGAACAGAAGGACTTTC 3′ and 5’ CCTGGGACCCTGACATAAAGTT using GoTaq^®^ qPCR Master Mix (Promega).

### Antibodies

Anti-PCM1 antibody was generated and used for immunofluorescence as previously described (Firat-Karalar et al, 2014). Other primary antibodies used for immunofluorescence in this study were rabbit anti-PCM1 (Proteintech) at 1:1000, goat anti-PCM1 (Santa Cruz Biotechnology) at 1:1000, mouse anti-γ-tubulin (GTU-88; Sigma-Aldrich) at 1:4000, mouse anti-GFP (3e6; Invitrogen) at 1:750, mouse anti-Myc (9e10; Sigma-Aldrich) at 1:500, mouse anti-acetylated tubulin (6-11-B; Sigma Aldrich) at 1:10.000, mouse anti-polyglutamylated tubulin (GT335, Adipogen) at 1:500, mouse anti-Smo (Santa Cruz Biotechnology) at 1:500, mouse anti-Arl13b (Neuromab) at 1:1000, rabbit anti-IFT88 (Proteintech) at 1:200, mouse anti-MIB1 (Santa Cruz Biotechnology) at 1:500, rabbit anti-CP110 (Proteintech) at 1:200, rabbit anti-Talpid3 (Proteintech) at 1:500, rabbit anti-Cep290 (Abcam) at 1:750, rat anti-ZO1 (Santa Cruz Biotechnology) at 1:500, rabbit anti-beta-catenin (Proteintech) at 1:500 and mouse anti-Cep164 (Santa Cruz Biotechnology) at 1:500. Secondary antibodies used for immunofluorescence experiments were AlexaFluor 488-, 568-or 633-coupled (Life Technologies) and they were used at 1:2000. Primary antibodies used for western blotting were rabbit anti-PCM1 (Proteintech) at 1:500, mouse anti-γ-tubulin (GTU-88; Sigma-Aldrich) at 1:4000, mouse anti-acetylated tubulin (6-11-B; Sigma Aldrich) at 1:10.000, mouse anti-polyglutamylated tubulin (GT335, Adipogen) at 1:500, mouse anti-Arl13b (Neuromab) at 1:500, mouse anti-Smo (Santa Cruz Biotechnology) at 1:500, rabbit anti-IFT88 (Proteintech) at 1:500, mouse anti-MIB1 (Sigma Aldrich) at 1:500, rabbit anti-CP110 (Proteintech) at 1:500 and rabbit anti-Cep164 (Proteintech) at 1:500, rabbit anti-Talpid3 (Proteintech) at 1:500, rabbit anti-Cep290 (Proteintech) at 1:500 rabbit, rabbit anti-BBS4 (Proteintech) at 1:500 and mouse anti-vinculin (Santa Cruz Biotechnology) at 1:1000. Secondary antibodies used for western blotting experiments were IRDye680- and IRDye 800-coupled and were used at 1:15000 (LI-COR Biosciences).

### Immunofluorescence, microscopy and quantitation

For immunofluorescence experiments, cells were grown on coverslips and fixed in either methanol or 4% PFA in PBS for indirect immunofluorescence. After rehydration in PBS, cells were blocked in 3% BSA (Sigma-Aldrich) in PBS + 0.1% Triton X-100. Coverslips were incubated in primary antibodies diluted in blocking solution and Alexa Fluor 488-, 594-, or 680-conjugated secondary antibodies (Invitrogen). Samples were mounted using Mowiol mounting medium containing N-propyl gallate (Sigma-Aldrich). Coverslips of cells were imaged using LAS X software (Premium; Leica) on a scanning confocal microscope (SP8; Leica Microsystems) with Plan Apofluar 63X 1.4 NA objective or a fluorescent microscope (DMi8; Leica Microsystems) with Plan Apofluar 63X 1.4 NA objective using a digital CMOS camera (Hamamatsu Orca Flash 4.0 V2 Camera). Images were processed using Photoshop (Adobe) or ImageJ (National Institutes of Health, Bethesda, MD).

Quantitative immunofluorescence of centrosomal and ciliary levels of proteins was performed on cells by acquiring a z-stack of control and depleted cells using identical gain and exposure settings, determined by adjusting settings based on the fluorescence signal in the control cells. The z-stacks were used to assemble maximum-intensity projections. The centrosome regions in these images were defined by γ-tubulin staining for each cell and the total pixel intensity of a round 3-μm^2^ area centered on the centrosome in each cell was measured using ImageJ and defined as the centrosomal intensity. The ciliary regions in these images were defined by Arl13b or acetylated tubulin for each cell. Background subtraction was performed by quantifying fluorescence intensity of a region of equal dimensions in the area neighboring the centrosome or cilium. Primary cilium formation was assessed by counting the total number of cells and the number of cells with primary cilia, as detected by Arl13b and acetylated tubulin staining. All data acquisition was done in a blinded manner.

### Transcriptome analysis

Transcriptome analysis was performed as described previously with the following modifications. Quality of reads of raw data was analyzed via FastQC program. We removed the sequencing reads with adaptor contamination and low-quality bases (quality score < Q30) via Trimmomatic (v0.35) to clean and advance the quality of the reads. All sequence data were 2×100 bp in length. The high quality reads were saved in fastq files and deposited to NCBI under accession number X. Clean and high quality data were utilized for the all downstream analysis. We used STAR aligner (v2.5.3) to generate genome indexes and mapping reads to the reference genomes that are hg38 for human cell line samples and GRCm38.p5 for mouse cell line samples. At the mapping step, reads having the non-canonical junctions were removed and only uniquely mapping reads were utilized. Reads Following the alignment, a Pearson’s correlation analysis was conducted to obtain the transcript-level R^2^ value between replicates. To determine the differentially expressed genes (DEGs) we utilized Cuffdiff with a minimum alignment count of 10, false discovery rate (FDR) < 0.05. The absolute value of |Fold change| ≥ 2 and q-value <0.05 was used as the threshold to judge significant differences in gene expression. DAVID Bioinformatics Resources 6.8 (Huang et al, 2008; Huang et al, 2009) was utilized to determine the statistical enrichment of the significantly (*p*-val < 0.05) differentially expressed genes in KEGG pathways and GO enrichment analysis. Analysis of GO annotation pathways provides information based on the biological processes (BP), cellular components (CC), and molecular functions (MF).

### Mass spectrometry

For the quantitative comparison of WT and knockout cells, snap frozen cell pellets of 1×10^6^ cells were resuspended in 50 µl PBS followed by the addition of 50 µl of 1% SDS in 100 mM Hepes, pH 8.4 and protease inhibitor cocktail (Roche, #11873580001). Samples were heated to 95°C for 5 min and then transferred on ice. Samples were treated with Benzonase (EMD Millipore Corp., #71206-3) at 37°C for 1 h to degrade DNA. Protein concentrations were determined and adjusted to 1 µg/µl. 20 µg thereof were subjected to an in-solution tryptic digest using a modified version of the Single-Pot Solid-Phase-enhanced Sample Preparation (SP3) protocol (Hughes et al, 2014; Moggridge et al, 2018). Here, lysates were added to Sera-Mag Beads (Thermo Scientific, #4515-2105-050250, 6515-2105-050250) in 10 µl 15% formic acid and 30 µl of ethanol. Binding of proteins was achieved by shaking for 15 min at room temperature. SDS was removed by 4 subsequent washes with 200 µl of 70% ethanol. Proteins were digested with 0.4 µg of sequencing grade modified trypsin (Promega, #V5111) in 40 µl Hepes/NaOH, pH 8.4 in the presence of 1.25 mM TCEP and 5 mM chloroacetamide (Sigma-Aldrich, #C0267) overnight at room temperature. Beads were separated, washed with 10 µl of an aqueous solution of 2% DMSO and the combined eluates were dried down. Peptides were reconstituted in 10 µl of H_2_O and reacted with 80 µg of TMT10plex (Thermo Scientific, #90111) label reagent dissolved in 4 µl of acetonitrile for 1 h at room temperature (Werner et al, 2014). Excess TMT reagent was quenched by the addition of 4 µl of an aqueous solution of 5% hydroxylamine (Sigma, 438227). Peptides were mixed to achieve a 1:1 ratio across all TMT-channels. Mixed peptides were subjected to a reverse phase clean-up step (OASIS HLB 96-well µElution Plate, Waters #186001828BA) and subjected to an off-line fractionation under high pH condition (Hughes et al, 2014). The resulting 12 fractions were then analyzed by LC-MS/MS on a Q Exactive Plus (Thermo Scientific) as previously described (PMID: 29706546). Briefly, peptides were separated using an UltiMate 3000 RSLC (Thermo Scientific) equipped with a trapping cartridge (Precolumn; C18 PepMap 100, 5 lm, 300 lm i.d. × 5 mm, 100 A°) and an analytical column (Waters nanoEase HSS C18 T3, 75 lm × 25 cm, 1.8 lm, 100 A°). Solvent A: aqueous 0.1% formic acid; Solvent B: 0.1% formic acid in acetonitrile (all solvents were of LC-MS grade). Peptides were loaded on the trapping cartridge using solvent A for 3 min with a flow of 30 µl/min. Peptides were separated on the analytical column with a constant flow of 0.3 µl/min applying a 2 h gradient of 2 – 28% of solvent B in A, followed by an increase to 40% B. Peptides were directly analyzed in positive ion mode applying with a spray voltage of 2.3 kV and a capillary temperature of 320°C using a Nanospray-Flex ion source and a Pico-Tip Emitter 360 lm OD × 20 lm ID; 10 lm tip (New Objective). MS spectra with a mass range of 375–1.200 m/z were acquired in profile mode using a resolution of 70.000 [maximum fill time of 250 ms or a maximum of 3e6 ions (automatic gain control, AGC)]. Fragmentation was triggered for the top 10 peaks with charge 2–4 on the MS scan (data-dependent acquisition) with a 30 second dynamic exclusion window (normalized collision energy was 32). Precursors were isolated with a 0.7m/z window and MS/MS spectra were acquired in profile mode with a resolution of 35,000 (maximum fill time of 120 ms or an AGC target of 2e5 ions).

Acquired data were analyzed using IsobarQuant (Franken et al, 2015) and Mascot V2.4 (Matrix Science) using a reverse UniProt FASTA Mus musculus database (UP000000589) including common contaminants. The following modifications were taken into account: Carbamidomethyl (C, fixed), TMT10plex (K, fixed), Acetyl (N-term, variable), Oxidation (M, variable) and TMT10plex (N-term, variable). The mass error tolerance for full scan MS spectra was set to 10 ppm and for MS/MS spectra to 0.02 Da. A maximum of 2 missed cleavages were allowed. A minimum of 2 unique peptides with a peptide length of at least seven amino acids and a false discovery rate below 0.01 were required on the peptide and protein level (Savitski et al, 2015).

### Mass Spectrometry Data Analysis

The proteins.txt output file of IsobarQuant was analyzed using the R programming language. As a quality control filter, we only allowed proteins that were quantified with at least two different unique peptides. The “signal_sum” columns were used and annotated to the knockout (KO) and wild-type (WT) conditions. Batch-effects were removed using the limma package (PMID: 25605792) and the resulting data were normalized with the vsn package (variance stabilization) (Hughes et al, 2014). Limma was employed again to look for differentially expressed proteins between KO and WT. We classified proteins as hits with a false discovery rate (FDR) smaller 5 % and a fold-change of at least 2-fold. Candidates were defined with an FDR smaller 20 % and a fold-change of at least 50%.

### Statistical Analysis

Statistical significance and p values were assessed by analysis of variance and Student's *t* tests using Prism software (GraphPad Software, La Jolla, CA). Error bars reflect SD. Following key is followed for asterisk placeholders for *p*-values in the figures: ****p<0.001 ***p<0.001, **p<0.01, *p<0.05.

## Acknowledgements

We acknowledge the Firat-Karalar lab members for insightful discussions regarding this work. In particular, we thank Melis Dilara Arslanhan for support during network analysis of the global proteomics data. IMCD3 cells were a kind gift from Moe Mahjoub (University of Washington, St. Louis). We thank the European Molecular Biology Laboratory (Heidelberg) for the use of the Proteomics Core Facility, which performed the TMT-based global proteome identification. This work was supported by ERC Grant 679140, Newton Advanced Fellowship RG84475 and TUBITAK Grant 214Z223 to ENF.

---

**Figure S1.**
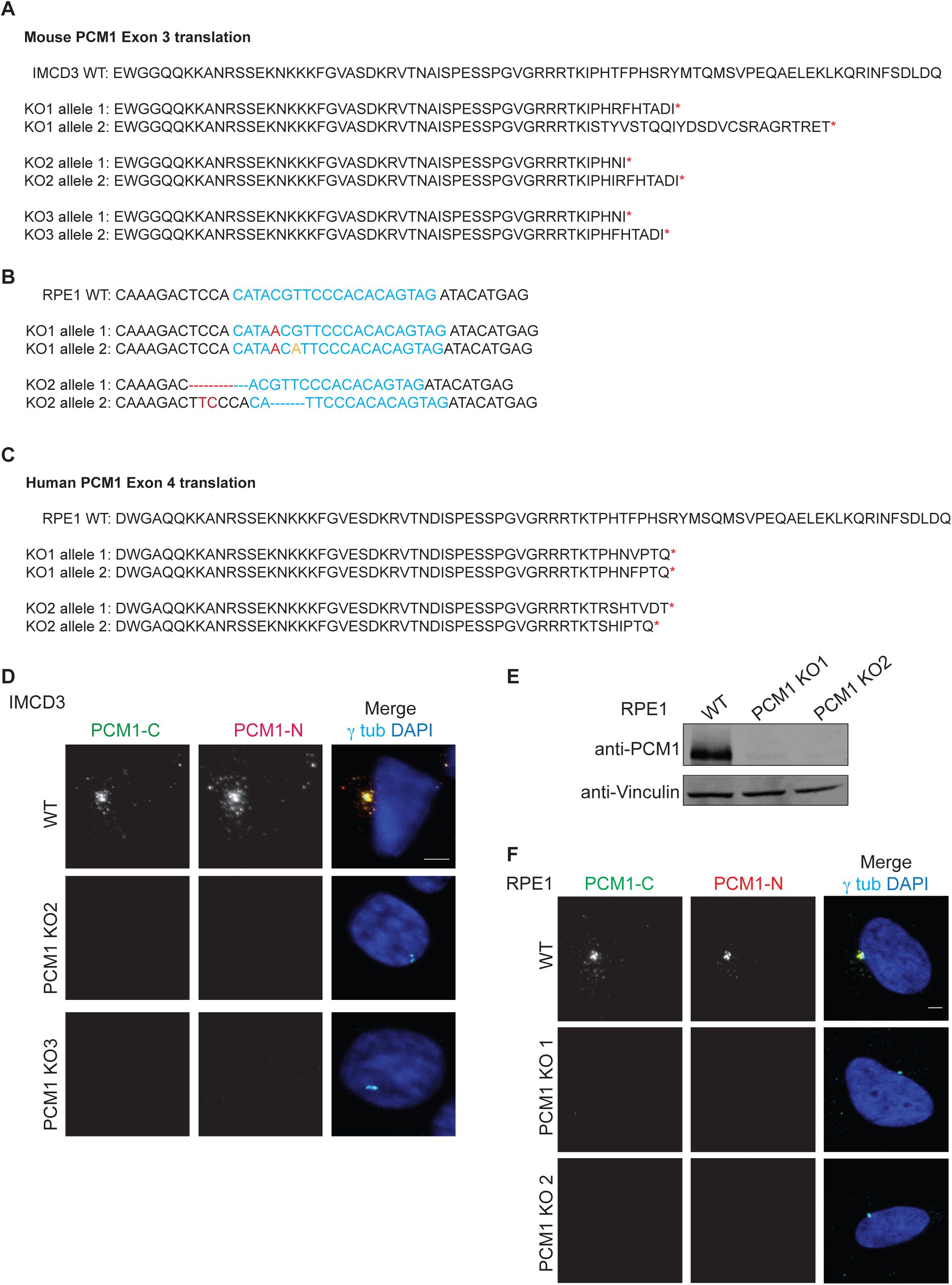
IMCD3 PCM1 KO and RPE1 PCM1 KO cells are devoid of satellite structures. (A) Translation products of the gRNA-targeting exon in IMCD3 KO clones. (B) RPE1 PCM1 KO clones are all compound heterozygotes with mutations that lead to early stop codons. 1000 bp region around the gRNA-target site was PCR amplified and cloned. Sequencing of five different clones for each line identified one 1nt insertion on one allele and one nt insertion and one nucleotide change on the other for line 1, 7 nt deletion on one allele and 4 nt deletion and and 2 nt insertion on the other for line 2. (C) Translation products of the gRNA-tareting exon in RPE1 PCM1 KO clones. (D) Immunofluorescence analysis of control and IMCD3 PCM1 KO clones. Cells were fixed and stained for centrosomes with anti-γ-tubulin antibody and PCM1 with PCM1-N antibody and PCM1-C antibody. (E, F) RPE1 PCM1 KO cells do not express PCM1. (D) Immunoblot analysis of whole-cell lysates from control cells and two RPE1 PCM1 KO clones with PCM1 antibody raised against its N-terminus. (E) Immunofluorescence analysis of control and RPE1 PCM1 KO clones. Cells were fixed and stained for centrosomes with anti-γ-tubulin antibody and PCM1 with PCM1-N antibody and PCM1-C antibody. Scale bar, 3 μm.

**Figure S2.**
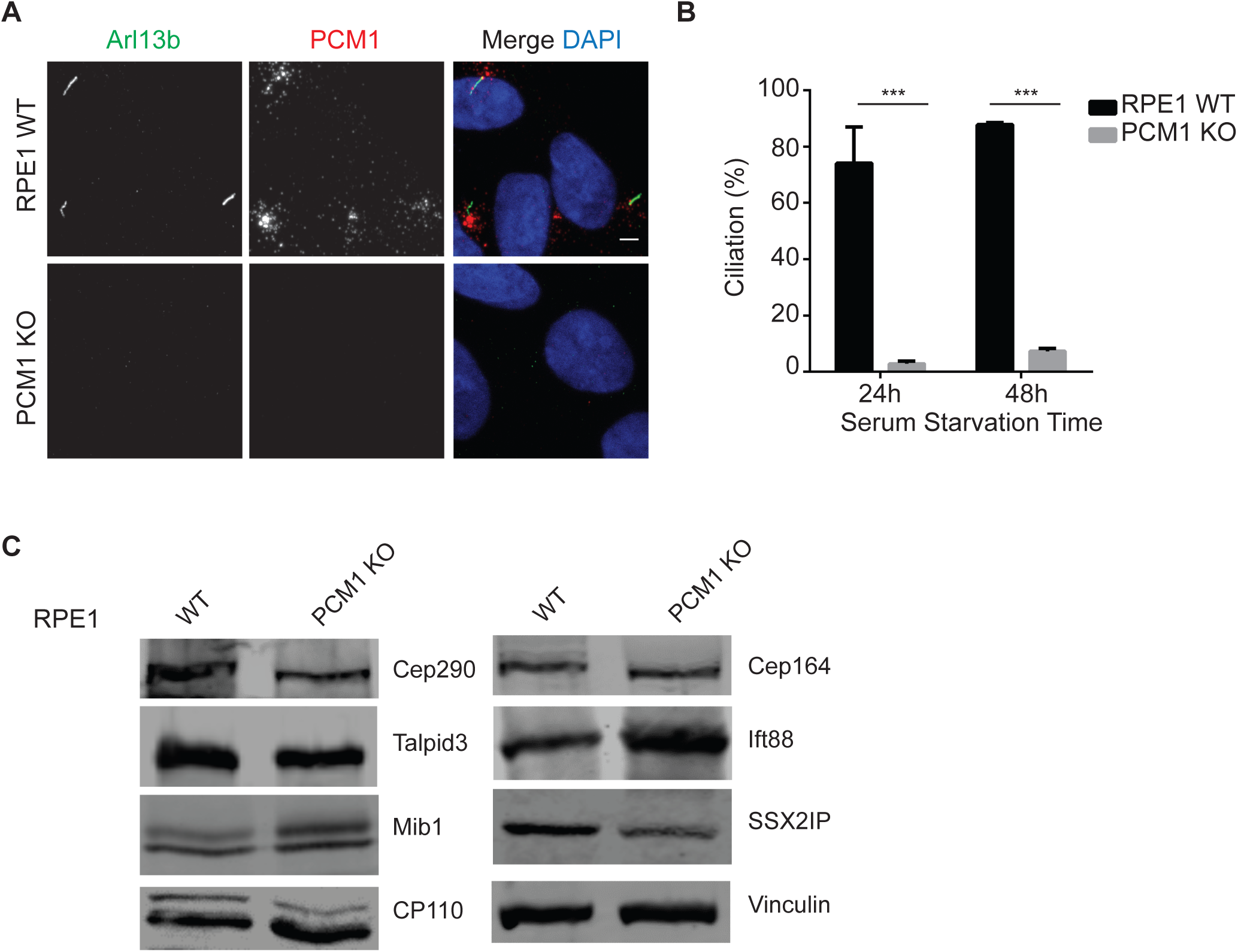
Satellites are essential for cilium assembly in RPE1 cells. (A) Effect of satellite loss on cilium formation. Control and RPE1 PCM1 KO cells were serum-starved for the indicated times and percentage of ciliated cells was determined by staining for acetylated-tubulin, Arl13b and DAPI. (B) Results shown are the mean of three independent experiments±SD (500 cells/experiment, ***p<0.001, t-test). (C) Effects of satellite loss on cellular abundance of centrosome proteins. Whole-cell lysates from control and RPE1 PCM1 KO cells were immunoblotted with the indicated antibodies. Vinculin was used as a loading control.

**Figure S3.**
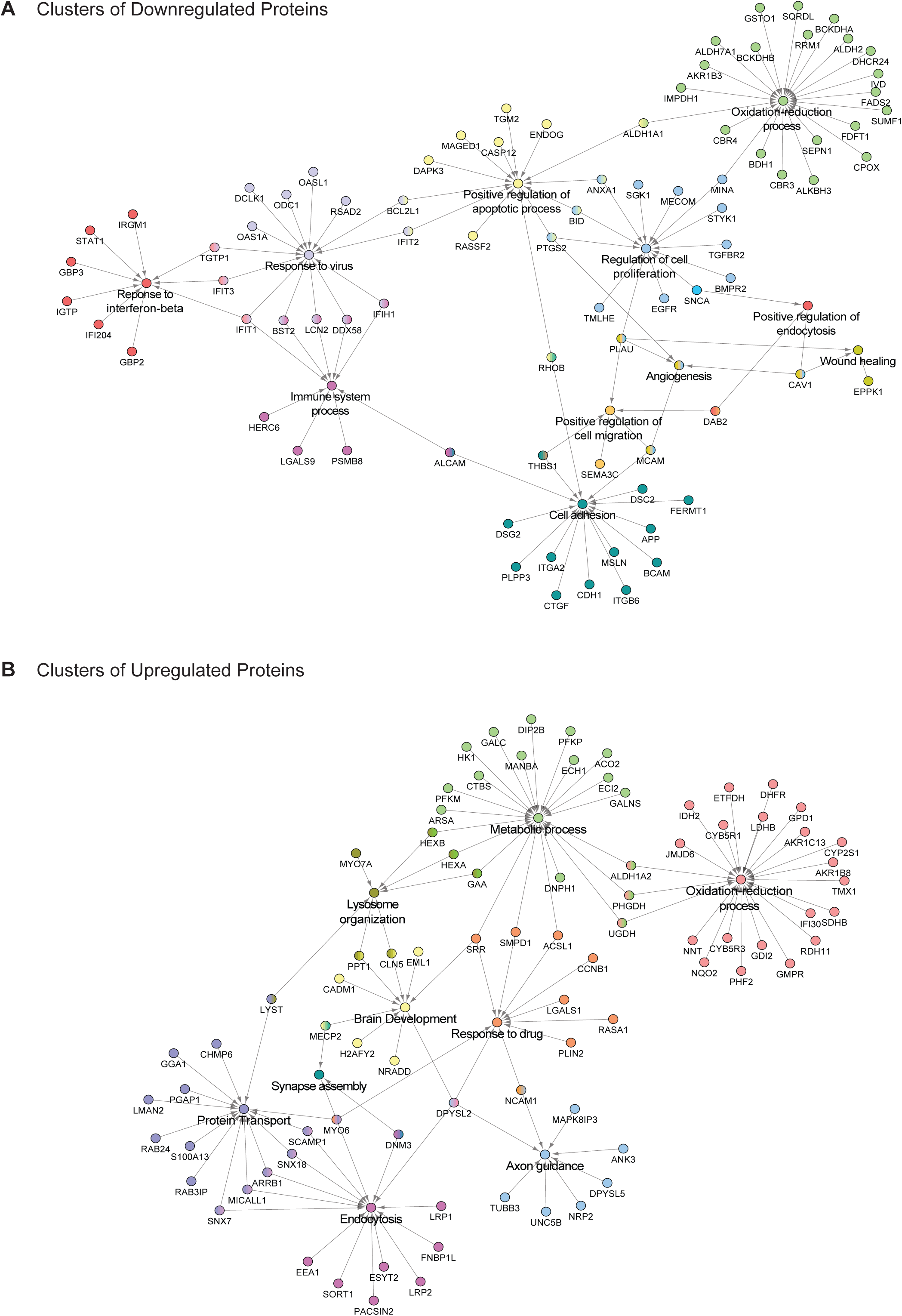
Interaction network analysis of the upregulated and downregulated proteins in IMCD3 PCM1 KO cells. Interaction network analysis for (A) downregulated and (B) upregulated proteins identified by GO-enrichment analysis revealed pathways enriched or depleted in satellite-less cells.

**Table S1.**
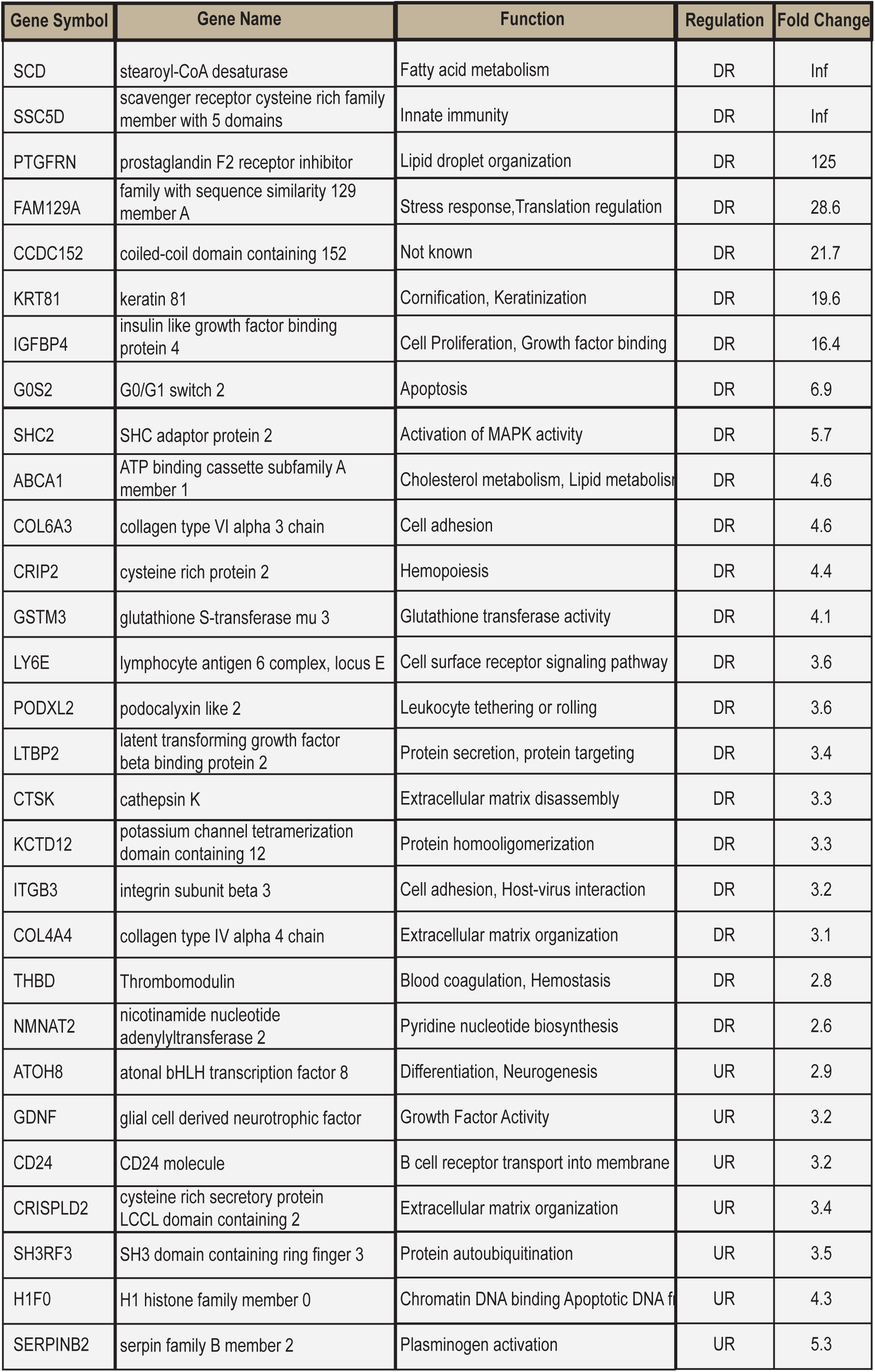
Differentially regulated genes in RPE1 PCM1 KO cells compared to control cells (> 1.5-fold, adj.P-value < 0.05) Genes significantly up- or down-regulated in RPE1 PCM1 KO cells compared to WT controls (>1.5-fol, adj. P-value<0.05)

**Table S2. List of all proteins identified in the TMT analysis of WT and IMCD3 PCM1 KO cells.**

**Table S3: List of all proteins analyzed by Limma and list of upregulated and downregulated proteins based on Limma analysis.** log_2_>0.5 for 295 upregulated proteins, log_2_<-0.5 for 296 downregulated proteins

## References

Akimov V, Rigbolt KT, Nielsen MM, Blagoev B (2011) Characterization of ubiquitination dependent dynamics in growth factor receptor signaling by quantitative proteomics. Mol Biosyst 7: 3223–3233

Barenz F, Mayilo D, Gruss OJ (2011) Centriolar satellites: busy orbits around the centrosome. Eur J Cell Biol 90: 983–989

Bauer M, Cubizolles F, Schmidt A, Nigg EA (2016) Quantitative analysis of human centrosome architecture by targeted proteomics and fluorescence imaging. EMBO J 35: 2152–2166

Bettencourt-Dias M, Hildebrandt F, Pellman D, Woods G, Godinho SA (2011) Centrosomes and cilia in human disease. Trends Genet 27: 307–315

Bhogaraju S, Cajanek L, Fort C, Blisnick T, Weber K, Taschner M, Mizuno N, Lamla S, Bastin P, Nigg EA, Lorentzen E (2013) Molecular basis of tubulin transport within the cilium by IFT74 and IFT81. Science 341: 1009–1012

Bradshaw NJ Porteous DJ (2012) DISC1-binding proteins in neural development, signalling and schizophrenia. Neuropharmacology 62: 1230–1241

Braun DA, Hildebrandt F (2017) Ciliopathies. Cold Spring Harbor perspectives in biology 9

Carvalho-Santos Z, Azimzadeh J, Pereira-Leal JB, Bettencourt-Dias M (2011) Evolution: Tracing the origins of centrioles, cilia, and flagella. J Cell Biol 194: 165–175

Cevik S, Hori Y, Kaplan OI, Kida K, Toivenon T, Foley-Fisher C, Cottell D, Katada T, Kontani K, Blacque OE (2010) Joubert syndrome Arl13b functions at ciliary membranes and stabilizes protein transport in Caenorhabditis elegans. J Cell Biol 188: 953–969

Corbit KC, Aanstad P, Singla V, Norman AR, Stainier DY, Reiter JF (2005) Vertebrate Smoothened functions at the primary cilium. Nature 437: 1018–1021

Dammermann A, Merdes A (2002) Assembly of centrosomal proteins and microtubule organization depends on PCM-1. J Cell Biol 159: 255–266

Datta SR, McQuillin A, Rizig M, Blaveri E, Thirumalai S, Kalsi G, Lawrence J, Bass NJ, Puri V, Choudhury K, Pimm J, Crombie C, Fraser G, Walker N, Curtis D, Zvelebil M, Pereira A, Kandaswamy R, St Clair D, Gurling HM (2010) A threonine to isoleucine missense mutation in the pericentriolar material 1 gene is strongly associated with schizophrenia. Mol Psychiatry 15: 615–628

Delous M, Hellman NE, Gaude HM, Silbermann F, Le Bivic A, Salomon R, Antignac C, Saunier S (2009) Nephrocystin-1 and nephrocystin-4 are required for epithelial morphogenesis and associate with PALS1/PATJ and Par6. Hum Mol Genet 18: 4711–4723

El-Brolosy MA, Stainier DYR (2017) Genetic compensation: A phenomenon in search of mechanisms. PLoS Genet 13: e1006780

Farina F, Gaillard J, Guerin C, Coute Y, Sillibourne J, Blanchoin L, Thery M (2016) The centrosome is an actin-organizing centre. Nat Cell Biol 18: 65–75

Firat-Karalar EN, Rauniyar N, Yates JR, 3rd, Stearns T (2014) Proximity interactions among centrosome components identify regulators of centriole duplication. Curr Biol 24: 664–670

Franken H, Mathieson T, Childs D, Sweetman GM, Werner T, Togel I, Doce C, Gade S, Bantscheff M, Drewes G, Reinhard FB, Huber W, Savitski MM (2015) Thermal proteome profiling for unbiased identification of direct and indirect drug targets using multiplexed quantitative mass spectrometry. Nat Protoc 10: 1567–1593

Ge X, Frank CL, Calderon de Anda F, Tsai LH (2010) Hook3 interacts with PCM1 to regulate pericentriolar material assembly and the timing of neurogenesis. Neuron 65: 191–203

Giles RH, Ajzenberg H, Jackson PK (2014) 3D spheroid model of mIMCD3 cells for studying ciliopathies and renal epithelial disorders. Nat Protoc 9: 2725–2731

Gupta GD, Coyaud E, Goncalves J, Mojarad BA, Liu Y, Wu Q, Gheiratmand L, Comartin D, Tkach JM, Cheung SW, Bashkurov M, Hasegan M, Knight JD, Lin ZY, Schueler M, Hildebrandt F, Moffat J, Gingras AC, Raught B, Pelletier L (2015) A Dynamic Protein Interaction Landscape of the Human Centrosome-Cilium Interface. Cell 163: 1484–1499

Gurling HM, Critchley H, Datta SR, McQuillin A, Blaveri E, Thirumalai S, Pimm J, Krasucki R, Kalsi G, Quested D, Lawrence J, Bass N, Choudhury K, Puri V, O'Daly O, Curtis D, Blackwood D, Muir W, Malhotra AK, Buchanan RW, Good CD, Frackowiak RS, Dolan RJ (2006) Genetic association and brain morphology studies and the chromosome 8p22 pericentriolar material 1 (PCM1) gene in susceptibility to schizophrenia. Arch Gen Psychiatry 63: 844–854

Hall EA, Keighren M, Ford MJ, Davey T, Jarman AP, Smith LB, Jackson IJ, Mill P (2013) Acute versus chronic loss of mammalian Azi1/Cep131 results in distinct ciliary phenotypes. PLoS Genet 9: e1003928

Haycraft CJ, Zhang Q, Song B, Jackson WS, Detloff PJ, Serra R, Yoder BK (2007) Intraflagellar transport is essential for endochondral bone formation. Development 134: 307–316

Hoang-Minh LB, Deleyrolle LP, Nakamura NS, Parker AK, Martuscello RT, Reynolds BA, Sarkisian MR (2016) PCM1 Depletion Inhibits Glioblastoma Cell Ciliogenesis and Increases Cell Death and Sensitivity to Temozolomide. Transl Oncol 9: 392–402

Hodges ME, Scheumann N, Wickstead B, Langdale JA, Gull K (2010) Reconstructing the evolutionary history of the centriole from protein components. J Cell Sci 123: 1407–1413

Hori A, Toda T (2017) Regulation of centriolar satellite integrity and its physiology. Cell Mol Life Sci 74: 213–229

Hu Q, Milenkovic L, Jin H, Scott MP, Nachury MV, Spiliotis ET, Nelson WJ (2010) A septin diffusion barrier at the base of the primary cilium maintains ciliary membrane protein distribution. Science 329: 436–439

Huang DW, Sherman BT, Lempicki RA (2008) Systematic and integrative analysis of large gene lists using DAVID bioinformatics resources. Nature Protocols 4: 44

Huang DW, Sherman BT, Lempicki RA (2009) Bioinformatics enrichment tools: paths toward the comprehensive functional analysis of large gene lists. Nucleic Acids Research 37: 1–13

Hughes CS, Foehr S, Garfield DA, Furlong EE, Steinmetz LM, Krijgsveld J (2014) Ultrasensitive proteome analysis using paramagnetic bead technology. Mol Syst Biol 10: 757

Huttlin EL, Bruckner RJ, Paulo JA, Cannon JR, Ting L, Baltier K, Colby G, Gebreab F, Gygi MP, Parzen H, Szpyt J, Tam S, Zarraga G, Pontano-Vaites L, Swarup S, White AE, Schweppe DK, Rad R, Erickson BK, Obar RA, Guruharsha KG, Li K, Artavanis-Tsakonas S, Gygi SP, Harper JW (2017) Architecture of the human interactome defines protein communities and disease networks. Nature 545: 505–509

Huttlin EL, Ting L, Bruckner RJ, Gebreab F, Gygi MP, Szpyt J, Tam S, Zarraga G, Colby G, Baltier K, Dong R, Guarani V, Vaites LP, Ordureau A, Rad R, Erickson BK, Wuhr M, Chick J, Zhai B, Kolippakkam D, Mintseris J, Obar RA, Harris T, Artavanis-Tsakonas S, Sowa ME, De Camilli P, Paulo JA, Harper JW, Gygi SP (2015) The BioPlex Network: A Systematic Exploration of the Human Interactome. Cell 162: 425–440

Insolera R, Shao W, Airik R, Hildebrandt F, Shi SH (2014) SDCCAG8 regulates pericentriolar material recruitment and neuronal migration in the developing cortex. Neuron 83: 805–822

Jakobsen L, Vanselow K, Skogs M, Toyoda Y, Lundberg E, Poser I, Falkenby LG, Bennetzen M, Westendorf J, Nigg EA, Uhlen M, Hyman AA, Andersen JS (2011) Novel asymmetrically localizing components of human centrosomes identified by complementary proteomics methods. EMBO J 30: 1520–1535

Joachim J, Razi M, Judith D, Wirth M, Calamita E, Encheva V, Dynlacht BD, Snijders AP, O'Reilly N, Jefferies HBJ, Tooze SA (2017) Centriolar Satellites Control GABARAP Ubiquitination and GABARAP-Mediated Autophagy. Curr Biol 27: 2123–2136 e2127

Kamiya A, Tan PL, Kubo K, Engelhard C, Ishizuka K, Kubo A, Tsukita S, Pulver AE, Nakajima K, Cascella NG, Katsanis N, Sawa A (2008) Recruitment of PCM1 to the centrosome by the cooperative action of DISC1 and BBS4: a candidate for psychiatric illnesses. Arch Gen Psychiatry 65: 996–1006

Kim J, Jo H, Hong H, Kim MH, Kim JM, Lee JK, Heo WD, Kim J (2015) Actin remodelling factors control ciliogenesis by regulating YAP/TAZ activity and vesicle trafficking. Nat Commun 6: 6781

Kim J, Krishnaswami SR, Gleeson JG (2008) CEP290 interacts with the centriolar satellite component PCM-1 and is required for Rab8 localization to the primary cilium. Hum Mol Genet 17: 3796–3805

Kim J, Lee JE, Heynen-Genel S, Suyama E, Ono K, Lee K, Ideker T, Aza-Blanc P, Gleeson JG (2010) Functional genomic screen for modulators of ciliogenesis and cilium length. Nature 464: 1048–1051

Kim K, Lee K, Rhee K (2012) CEP90 is required for the assembly and centrosomal accumulation of centriolar satellites, which is essential for primary cilia formation. PLoS One 7: e48196

Kodani A, Yu TW, Johnson JR, Jayaraman D, Johnson TL, Al-Gazali L, Sztriha L, Partlow JN, Kim H, Krup AL, Dammermann A, Krogan NJ, Walsh CA, Reiter JF (2015) Centriolar satellites assemble centrosomal microcephaly proteins to recruit CDK2 and promote centriole duplication. eLife 4

Kohli P, Hohne M, Jungst C, Bertsch S, Ebert LK, Schauss AC, Benzing T, Rinschen MM, Schermer B (2017) The ciliary membrane-associated proteome reveals actin-binding proteins as key components of cilia. EMBO Rep 18: 1521–1535

Kubo A, Sasaki H, Yuba-Kubo A, Tsukita S, Shiina N (1999) Centriolar satellites: molecular characterization, ATP-dependent movement toward centrioles and possible involvement in ciliogenesis. J Cell Biol 147: 969–980

Kubo A, Tsukita S (2003) Non-membranous granular organelle consisting of PCM-1: subcellular distribution and cell-cycle-dependent assembly/disassembly. J Cell Sci 116: 919–928

Lopes CA, Prosser SL, Romio L, Hirst RA, O'Callaghan C, Woolf AS, Fry AM (2011) Centriolar satellites are assembly points for proteins implicated in human ciliopathies, including oral-facial-digital syndrome 1. J Cell Sci 124: 600–612

Luders J, Stearns T (2007) Microtubule-organizing centres: a re-evaluation. Nat Rev Mol Cell Biol 8: 161–167

Mahjoub MR, Stearns T (2012) Supernumerary centrosomes nucleate extra cilia and compromise primary cilium signaling. Curr Biol 22: 1628–1634

Mahjoub MR, Xie Z, Stearns T (2010) Cep120 is asymmetrically localized to the daughter centriole and is essential for centriole assembly. J Cell Biol 191: 331–346

Mirvis M, Stearns T, James Nelson W (2018) Cilium structure, assembly, and disassembly regulated by the cytoskeleton. Biochem J 475: 2329–2353

Moggridge S, Sorensen PH, Morin GB, Hughes CS (2018) Extending the Compatibility of the SP3 Paramagnetic Bead Processing Approach for Proteomics. J Proteome Res 17: 1730–1740

Nachury MV (2014) How do cilia organize signalling cascades? Philos Trans R Soc Lond B Biol Sci 369

Nachury MV, Loktev AV, Zhang Q, Westlake CJ, Peranen J, Merdes A, Slusarski DC, Scheller RH, Bazan JF, Sheffield VC, Jackson PK (2007) A core complex of BBS proteins cooperates with the GTPase Rab8 to promote ciliary membrane biogenesis. Cell 129: 1201–1213

Nielsen JC, Nordgaard C, Tollenaere MAX, Bekker-Jensen S (2018) Osmotic Stress Blocks Mobility and Dynamic Regulation of Centriolar Satellites. Cells 7

Nigg EA, Cajanek L, Arquint C (2014) The centrosome duplication cycle in health and disease. FEBS Lett 588: 2366–2372

Nigg EA, Holland AJ (2018) Once and only once: mechanisms of centriole duplication and their deregulation in disease. Nat Rev Mol Cell Biol 19: 297–312

Obino D, Farina F, Malbec O, Saez PJ, Maurin M, Gaillard J, Dingli F, Loew D, Gautreau A, Yuseff MI, Blanchoin L, Thery M, Lennon-Dumenil AM (2016) Actin nucleation at the centrosome controls lymphocyte polarity. Nat Commun 7: 10969

Otto EA, Hurd TW, Airik R, Chaki M, Zhou W, Stoetzel C, Patil SB, Levy S, Ghosh AK, Murga-Zamalloa CA, van Reeuwijk J, Letteboer SJ, Sang L, Giles RH, Liu Q, Coene KL, Estrada-Cuzcano A, Collin RW, McLaughlin HM, Held S, Kasanuki JM, Ramaswami G, Conte J, Lopez I, Washburn J, Macdonald J, Hu J, Yamashita Y, Maher ER, Guay-Woodford LM, Neumann HP, Obermuller N, Koenekoop RK, Bergmann C, Bei X, Lewis RA, Katsanis N, Lopes V, Williams DS, Lyons RH, Dang CV, Brito DA, Dias MB, Zhang X, Cavalcoli JD, Nurnberg G, Nurnberg P, Pierce EA, Jackson PK, Antignac C, Saunier S, Roepman R, Dollfus H, Khanna H, Hildebrandt F (2010) Candidate exome capture identifies mutation of SDCCAG8 as the cause of a retinal-renal ciliopathy. Nat Genet 42: 840–850

Pampliega O, Orhon I, Patel B, Sridhar S, Diaz-Carretero A, Beau I, Codogno P, Satir BH, Satir P, Cuervo AM (2013) Functional interaction between autophagy and ciliogenesis. Nature 502: 194–200

Paz J, Luders J (2018) Microtubule-Organizing Centers: Towards a Minimal Parts List. Trends Cell Biol 28: 176–187

Pazour GJ, Baker SA, Deane JA, Cole DG, Dickert BL, Rosenbaum JL, Witman GB, Besharse JC (2002) The intraflagellar transport protein, IFT88, is essential for vertebrate photoreceptor assembly and maintenance. J Cell Biol 157: 103–113

Pazour GJ, Dickert BL, Vucica Y, Seeley ES, Rosenbaum JL, Witman GB, Cole DG (2000) Chlamydomonas IFT88 and its mouse homologue, polycystic kidney disease gene tg737, are required for assembly of cilia and flagella. J Cell Biol 151: 709–718

Rai AK, Chen JX, Selbach M, Pelkmans L (2018) Kinase-controlled phase transition of membraneless organelles in mitosis. Nature 559: 211–216

Reiter JF, Leroux MR (2017) Genes and molecular pathways underpinning ciliopathies. Nat Rev Mol Cell Biol 18: 533–547

Rossi A, Kontarakis Z, Gerri C, Nolte H, Holper S, Kruger M, Stainier DY (2015) Genetic compensation induced by deleterious mutations but not gene knockdowns. Nature 524: 230–233

Sadrian B, Cheng TW, Shull O, Gong Q (2012) Rap1gap2 regulates axon outgrowth in olfactory sensory neurons. Mol Cell Neurosci 50: 272–282

Savitski MM, Wilhelm M, Hahne H, Kuster B, Bantscheff M (2015) A Scalable Approach for Protein False Discovery Rate Estimation in Large Proteomic Data Sets. Mol Cell Proteomics 14: 2394–2404

Schmidt TI, Kleylein-Sohn J, Westendorf J, Le Clech M, Lavoie SB, Stierhof YD, Nigg EA (2009) Control of centriole length by CPAP and CP110. Curr Biol 19: 1005–1011

Sha J, Han Q, Chi C, Zhu Y, Pan J, Dong B, Huang Y, Xia W, Xue W (2018) PRKAR2B promotes prostate cancer metastasis by activating Wnt/beta-catenin and inducing epithelial-mesenchymal transition. J Cell Biochem 119: 7319–7327

Sha J, Xue W, Dong B, Pan J, Wu X, Li D, Liu D, Huang Y (2017) PRKAR2B plays an oncogenic role in the castration-resistant prostate cancer. Oncotarget 8: 6114–6129

Sorokin S (1962) Centrioles and the formation of rudimentary cilia by fibroblasts and smooth muscle cells. J Cell Biol 15: 363–377

Sorokin SP (1968) Reconstructions of centriole formation and ciliogenesis in mammalian lungs. J Cell Sci 3: 207–230

Srsen V, Fant X, Heald R, Rabouille C, Merdes A (2009) Centrosome proteins form an insoluble perinuclear matrix during muscle cell differentiation. BMC Cell Biol 10: 28

Srsen V, Gnadt N, Dammermann A, Merdes A (2006) Inhibition of centrosome protein assembly leads to p53-dependent exit from the cell cycle. J Cell Biol 174: 625–630

Staples CJ, Myers KN, Beveridge RD, Patil AA, Howard AE, Barone G, Lee AJ, Swanton C, Howell M, Maslen S, Skehel JM, Boulton SJ, Collis SJ (2014) Ccdc13 is a novel human centriolar satellite protein required for ciliogenesis and genome stability. J Cell Sci 127: 2910–2919

Staples CJ, Myers KN, Beveridge RD, Patil AA, Lee AJ, Swanton C, Howell M, Boulton SJ, Collis SJ (2012) The centriolar satellite protein Cep131 is important for genome stability. J Cell Sci 125: 4770–4779

Stowe TR, Wilkinson CJ, Iqbal A, Stearns T (2012) The centriolar satellite proteins Cep72 and Cep290 interact and are required for recruitment of BBS proteins to the cilium. Mol Biol Cell 23: 3322–3335

Tabares-Seisdedos R, Rubenstein JL (2009) Chromosome 8p as a potential hub for developmental neuropsychiatric disorders: implications for schizophrenia, autism and cancer. Mol Psychiatry 14: 563–589

Tang Z, Lin MG, Stowe TR, Chen S, Zhu M, Stearns T, Franco B, Zhong Q (2013) Autophagy promotes primary ciliogenesis by removing OFD1 from centriolar satellites. Nature 502: 254–257

Tollenaere MA, Mailand N, Bekker-Jensen S (2015) Centriolar satellites: key mediators of centrosome functions. Cell Mol Life Sci 72: 11–23

Villumsen BH, Danielsen JR, Povlsen L, Sylvestersen KB, Merdes A, Beli P, Yang YG, Choudhary C, Nielsen ML, Mailand N, Bekker-Jensen S (2013) A new cellular stress response that triggers centriolar satellite reorganization and ciliogenesis. EMBO J 32: 3029–3040

Vladar EK, Stearns T (2007) Molecular characterization of centriole assembly in ciliated epithelial cells. J Cell Biol 178: 31–42

Wang L, Dynlacht BD (2018) The regulation of cilium assembly and disassembly in development and disease. Development 145

Wang L, Lee K, Malonis R, Sanchez I, Dynlacht BD (2016) Tethering of an E3 ligase by PCM1 regulates the abundance of centrosomal KIAA0586/Talpid3 and promotes ciliogenesis. eLife 5

Werner T, Sweetman G, Savitski MF, Mathieson T, Bantscheff M, Savitski MM (2014) Ion coalescence of neutron encoded TMT 10-plex reporter ions. Anal Chem 86: 3594–3601

Wu CT, Chen HY, Tang TK (2018) Myosin-Va is required for preciliary vesicle transportation to the mother centriole during ciliogenesis. Nat Cell Biol 20: 175–185

Zhang W, Kim PJ, Chen Z, Lokman H, Qiu L, Zhang K, Rozen SG, Tan EK, Je HS, Zeng L (2016) MiRNA-128 regulates the proliferation and neurogenesis of neural precursors by targeting PCM1 in the developing cortex. eLife 5

